# Fungi are more dispersal limited than bacteria among flowers

**DOI:** 10.1101/2020.05.19.104968

**Authors:** Rachel L. Vannette, Marshall S. McMunn, Griffin W. Hall, Tobias G. Mueller, Ivan Munkres, Douglas Perry

## Abstract

Variation in dispersal ability among taxa affects community assembly and biodiversity maintenance within metacommunities. Although fungi and bacteria frequently coexist, their relative dispersal abilities are poorly understood. Nectar-inhabiting microbial communities affect plant reproduction and pollinator behavior, and are excellent models for studying dispersal of bacteria and fungi in a metacommunity framework. Here, we assay dispersal ability of common nectar bacteria and fungi in an insect-based dispersal experiment. We then compare these results to the incidence and abundance of culturable flower-inhabiting bacteria and fungi within naturally occurring flowers across two coflowering communities in California across two flowering seasons. Our microbial dispersal experiment demonstrates that bacteria disperse among habitat patches more readily than fungi via thrips. Across all flowers, bacterial and fungal incidence and abundance were positively correlated but bacteria were much more widespread, suggesting shared dispersal routes or habitat requirements but differences in dispersal and colonization frequency. The finding that bacteria are more common among flowers sampled here, in part due to superior insect-mediated dispersal, may have broad relevance for microbial life-history, community assembly of microbes and plant-pollinator interactions.

## Introduction

Variation in dispersal ability among species can affect the composition and function of individual communities and biodiversity maintenance within metacommunities (Mouquet and Loreau 2003, Leibold et al. 2004). Dispersal limitation is well-documented for macro-organisms, and increasing evidence suggests that microbial taxa including fungi and bacteria can also be dispersal limited (Lindow and Andersen 1996, Peay et al. 2012, Albright and Martiny 2018, Svoboda et al. 2018). Despite the fact that bacteria and fungi often co-occur within communities, few studies have examined their relative dispersal abilities (but see Barberán et al. 2015). This gap in understanding is likely due to challenges in microbial detection, species delimitation using amplicon sequencing, and limited natural history knowledge of many microorganisms (Kraemer and Boynton 2016), particularly in a community context. As a result, comparisons in dispersal ability between coexisting bacteria and fungi are rare and limit the ability to predict the relative importance of dispersal for microbial community ecology and assembly dynamics. In addition, it is likely that processes underlying microbial community assembly and dynamics differ from those structuring communities of macroorganisms (Koskella et al. 2017). Characterizing dispersal differences of microbial taxa under realistic conditions is important to understanding how dispersal may shape microbial community assembly.

The microbial community in floral nectar—a sugar-rich reward that plants provide to pollinators--is ecologically important and a tractable model system for comparing bacterial and fungal dispersal limitation (Chappell and Fukami 2018, Vannette 2020). The composition of bacteria and fungi in flowers can influence plant reproduction (Herrera et al. 2013, Schaeffer and Irwin 2014), either directly (Alexander and Antonovics 1988, Ngugi and Scherm 2006) or indirectly through effects on pollinator behavior and health (Vannette et al. 2013, Lenaerts et al. 2017, Sobhy et al. 2018). Bacteria and fungi often co-occur in nectar (Herrera et al. 2009, Fridman et al. 2012) where they can attain high abundance (Fridman et al. 2012, Tsuji and Fukami 2018). The bacteria and fungi that grow in nectar are a small subset of the microbial communities introduced to nectar (Herrera et al. 2010, Belisle et al. 2012, Pozo et al. 2012, Alvarez-Pérez and Herrera 2013) and exhibit traits that may enhance fitness in nectar environments (Álvarez-Pérez et al. 2012, Pozo et al. 2012, Dhami et al. 2016a).

Despite strong initial environmental filtering, nectar-inhabiting microbial communities are ideal for examining dispersal limitation for a few reasons. First, communities in nectar undergo primary succession (Chappell and Fukami 2018). Few culturable microbes are found in nectar of newly opened flowers that have not received visitation by animals (Brysch-Herzberg 2004b, Vannette and Fukami 2017, von Arx et al. 2019, Morris et al. 2020), but flower visitors introduce bacteria and fungi that successfully colonize floral nectar (Lachance et al. 2001, Herrera et al. 2008). Second, the majority of bacteria and fungi that survive and thrive in floral nectar are culturable (Morris et al. 2020)— shotgun sequencing of nectar microbial communities reveal qualitatively similar taxonomic composition to culture-based analyses. Third, bacteria and fungi in nectar are thought to depend on phoresy via flower visitors (Brysch-Herzberg 2004b, Vannette and Fukami 2017, Morris et al. 2020). Fourth, plant species differ in flower morphologies associated with variation in nectar accessibility to wind and animal pollinators.

In addition, the assembly processes shaping nectar microbial communities are thought to depend on differential dispersal limitation. Laboratory experiments and field observations suggest that inhibitory priority effects (interactions in which species arrival order determines their outcome) influence community structure in floral nectar (Tucker and Fukami 2014, Toju et al. 2018) via differential dispersal of species to flowers (Mittelbach et al. 2016). However, positive interactions among nectar-inhabiting taxa have been suggested (Alvarez-Pérez and Herrera 2013), so the relative influence of dispersal and species interactions (including competition) on nectar microbe metacommunities are not clear. Elucidating patterns of bacteria and fungi dispersal limitation and co-occurrence in nectar would shed light on the mechanisms underlying their community assembly, inform the competitive environments experienced by bacteria and fungi in nectar, and have implications for microbial effects on interactions between plants and floral visitors.

Here, we test the hypothesis that nectar-inhabiting bacteria and fungi differ in dispersal and occupancy among individual flowers in two coflowering communities. We assess realized dispersal—which integrates the components of emigration (leaving a habitat), survival during transport, and establishment in a new habitat (flower)—which ultimately determines community membership (Baguette et al. 2012). First, we examined microbial dispersal ability empirically and second we assessed bacterial and fungal occupancy (incidence) and abundance in individual nectar samples of more than 1800 flowers, spanning 43 species of plants at two sites to examine 1) do bacteria or fungi differ in dispersal ability by a common flower-visiting insect; 2) are patterns of bacterial and fungal incidence or abundance in nectar similar across a plant community; 3) does microbial presence or abundance differ among plant species or with floral accessibility (flower morphology); 4) does season, year or site predict patterns of bacterial or fungal incidence and abundance?

## Materials and methods

### Insect-based microbial dispersal assay

To empirically test whether nectar microbes differ in their dispersal ability, we developed an assay of microbial dispersal resulting from the feeding activity of western flower thrips (*Frankliniella occidentalis*). The assay consisted of 5 sterilized adult female thrips freely foraging within a 96-well plate in which 1/3 of wells contained a microbial suspension (200uL 10,000 cells/uL) in sweetened tryptic soy broth (TSB, adding 15% sucrose and 15% fructose) and the remaining 2/3 of wells were filled with sterile TSB (200uL). Thrips were sourced from a colony held by Diane Ullman (see Supplementary Figure, Methods S1 for assay methods details). Microbial isolates from field sampling in this study (see below and Supplementary Table S1), and other nectar-associated microbe collections were used (6 bacteria, 5 fungi). After thrips foraged for 24 hours at 30°C under 16L:8D light, thrips were killed by adding 320uL of ethyl acetate to the plate bottom and the media was incubated for 5 additional days. Control replicates with thrips and no added microbes were included in all trials to ensure adequate sterilization and trial consistency. Following incubation, we assessed microbial occupancy of initially sterile wells by calculating the deviation of optical density (OD 600nm) from a blank control plate (no thrips, no microbes) with an occupancy threshold equal to the mean OD of control wells +6 standard deviations.

### Coflowering community sampling for bacteria and fungi

To assess patterns of microbial incidence and abundance, we sampled standing crop floral nectar during 2016 and 2017 at two sites in northern California: Stebbins Cold Canyon Reserve (38.49806°, −122.09931° 140m, hereafter Stebbins) in Winters, CA and at flowering plots maintained at the Laidlaw Honey Bee Facility (38.53653°, − 121.78864°, 19m, hereafter Bee Biology) in Davis, CA, which hosts an active apiary. The sites are approximately 30 kilometers apart, and differ in pollinator species composition and anthropogenic influence, but share a subset of plant species.

Between late February and early July, plant communities were sampled every two to four weeks. We collected approximately 10 flowers from abundant plant species. In some cases, collections were limited by floral availability and reserve collection restrictions to protect plant populations. When possible, flowers were sampled from multiple individual plants and sub-populations or plots (Supplementary Table S2). Care was taken to sample flowers that had been open more than one day when possible, to allow the opportunity for floral visitation and microbial immigration to flowers. Individual inflorescences were collected, placed upright in humidified boxes and kept cool until nectar extraction and plating, no more than 5 hours later. In the lab, flowers were destructively sampled. Nectar was collected using 10 μl microcapillary tubes, and its volume was quantified. Nectar was diluted in 30 μl of sterile water (D0), then diluted 10 and 100 fold (D1 and D2 respectively) in sterile phosphate-buffered saline (Dulbecco’s, pH 7.1-7.5). To assess fungal and bacterial abundance, 50 μl of the 10 fold dilution (containing 5 μl of D0) and 50 μl of the 100 fold dilution (containing 0.5μl of D0) were plated on yeast media agar (YMA) containing the antibacterial chloramphenicol and Reasoner’s agar containing the antifungal cycloheximide (R2A Oxoid formula with 16% sucrose), respectively. All samples were plated the day of collection. For convenience, we refer to the total number of colonies on YMA as “fungi” and colonies on R2A as “bacteria” throughout the manuscript although some colonies on each media type may be comprised by microbes resistant to the antimicrobial compounds used here (e.g. bacteria resistant to chloramphenicol (Dhami et al. 2018)). Based on dilution curves, we were able to detect CFUs on plates at approximately 10^1^ live cells in the original nectar sample for YMA and 10^2^ cells for R2A (Supplementary methods S2 and Figure S2).

Negative controls were plated for each sampling bout to detect potential contamination and samples were discarded if contamination was detected (detected for 1 collection date; these samples were removed from the analysis). Agar plates were incubated at 28°C and colony-forming units (CFUs) were counted after 48-72 hours. The total number of CFUs and CFU density for each nectar sample was calculated based on dilutions and original sample volume. Over the course of the study, 1,825 nectar samples were collected and plated on two media types. In 2016, colonies were picked haphazardly from plates collected over the entire season including all sites and plant species and frozen in glycerol.

### Nectar accessibility: flower corolla length

Microbial dispersal to nectar is likely affected by the location and accessibility of floral nectar within the flower. Specifically, nectar that is not enclosed by flower tissue may be more easily accessed via wind dispersal and may also be accessible to a greater diversity of insect visitors. On the other hand, nectar protected by flower tissues comprising the perianth (non-reproductive structures of the flower) may be less accessible via wind dispersal and contacted only when flowers are visited by long-tongued birds or insects (Kingston and Mc Quillan 2000, Lara and Ornelas 2001). We examined if bacterial or fungal incidence or abundance depended on nectar location within the flower (flower corolla length). Nectar within flowers of each species were classified as “exposed” (no corolla/exposed nectary), or contained within “short” (1-5 mm), “mid” (5-15 mm), or “long” (15+ mm) perianth, based on corolla tube length (Supplementary table S2).

### Microbial identification: culture-based and culture-independent analyses

A subset of microbial strains from glycerol stocks were identified using MALDI-TOF and spectra were compared to Bruker Bacteria and Eukaryote libraries and a custom inhouse database curated from previously identified microbial isolates from nectar (Supplementary methods S3). In addition, we used a culture-independent sequencing approach to characterize diversity in full-length 16S and ITS sequences from 25 pooled nectar samples (239 flowers in each sample), representing multiple sampling dates at each site from 2016. Pooled nectar samples were extracted using the Nucleospin Microbial DNA kit (Macherey-Nagel, Bethlehem, PA, USA) and submitted for full-length PacBio16S and ITS sequencing performed at Dalhousie IMR. Recovered sequences were processed using dada2 pipeline for full-length rRNA genes (Callahan et al. 2016, Callahan et al. 2019) for recovery of actual sequence variants (ASVs). Briefly, primer sequences were removed, and resulting sequences were quality filtered, error-corrected and denoised followed by taxonomy assignment using SILVA v. 138 training set (Quast et al. 2012) for bacteria and UNITE 4/2/2020 release for fungi (Kõljalg et al. 2005) followed by chimera detection and removal (See Dryad https://doi.org/10.25338/B8403V for bioinformatics scripts). ASVs annotated as chloroplast or plant, and those that were also obtained from extraction blanks were removed.

### Statistical analysis

To examine if microbes differed in dispersal ability in our experimental setup, the proportion of wells colonized by a microbe within a plate (calculated using OD as described above; plates as replicates) was analyzed using a binomial GLMM (package lme4) to compare bacterial and fungal dispersal plates with strain as a random effect. A t-test was also used to compare bacteria and fungi by the mean proportion of wells colonized by each strain across 8 replicate plates.

To examine the relationship between bacterial and fungal incidence and abundance in individual nectar samples, we used Pearson correlations. In addition, we report directly calculated conditional probabilities and a χ^2^ test to test for non-independence of bacterial and fungal incidence amongst flowers.

To compare plant species differed in bacterial or fungal presence or abundance, we used logistic regression and linear regression with plant species as a predictor. To compare if nectar accessibility quantified using corolla length category was associated with bacterial or fungal presence or abundance, we used logistic regression and linear regression with nectary location (exposed, short corolla, mid corolla or long corolla) as a predictor.

To determine if bacteria and fungi differ in incidence in floral nectar with Julian date, between sites, or between years, we used a logistic regression implemented using ‘glm’, using bacteria or fungal presence as a response and site, year, and plant species, and their two-way interactions, as predictors. We compared among: models containing all interactions, models with non-significant interactions removed, and models with no interactions using AIC, and from the best-fit model, significance of each term was assessed using likelihood ratio tests. To determine if bacteria or fungi differ in abundance in floral nectar across years or sites, or with Julian date, we used a linear regression implemented using ‘lm’, with log_10_(CFUs+1) as a response and site, Julian date, and year as predictors. Model selection was performed on bacteria and fungi separately using AIC as described above, and significance was assessed using likelihood ratio tests.

We assessed the frequency with which microbial genera were detected using MALDI-TOF among flower morphologies, or across the season (early, mid, or late, see supplement for details) using a χ^2^ test for contingency tables. Microbial composition from PacBio sequencing is presented to compare taxonomic composition with MALDI-TOF data but due to limited numbers of samples successfully sequenced, no formal analysis was performed. All analyses were conducted using RStudio and R v. 3.6.3 (Johnson et al. 2009).

## Results

### Insect-based dispersal assay

Fungi were less able to disperse among wells than bacteria in the thrips dispersal assay (Figure 1). Using strain-level averages to compare dispersal abilities between kingdoms, bacteria colonized 31% more wells than did fungi over 24 hours of thrips activity (t-test, strains as replicates, t = 4.31, p=0.002). Results from the GLMM confirm this difference between fungi and bacteria average dispersal ability (z-value =-4.66, p<0.001). Control treatments had detectable, but low levels of contamination (higher than two fungal strains) despite exhaustive measures to sterilize thrips.

**Figure 1.**
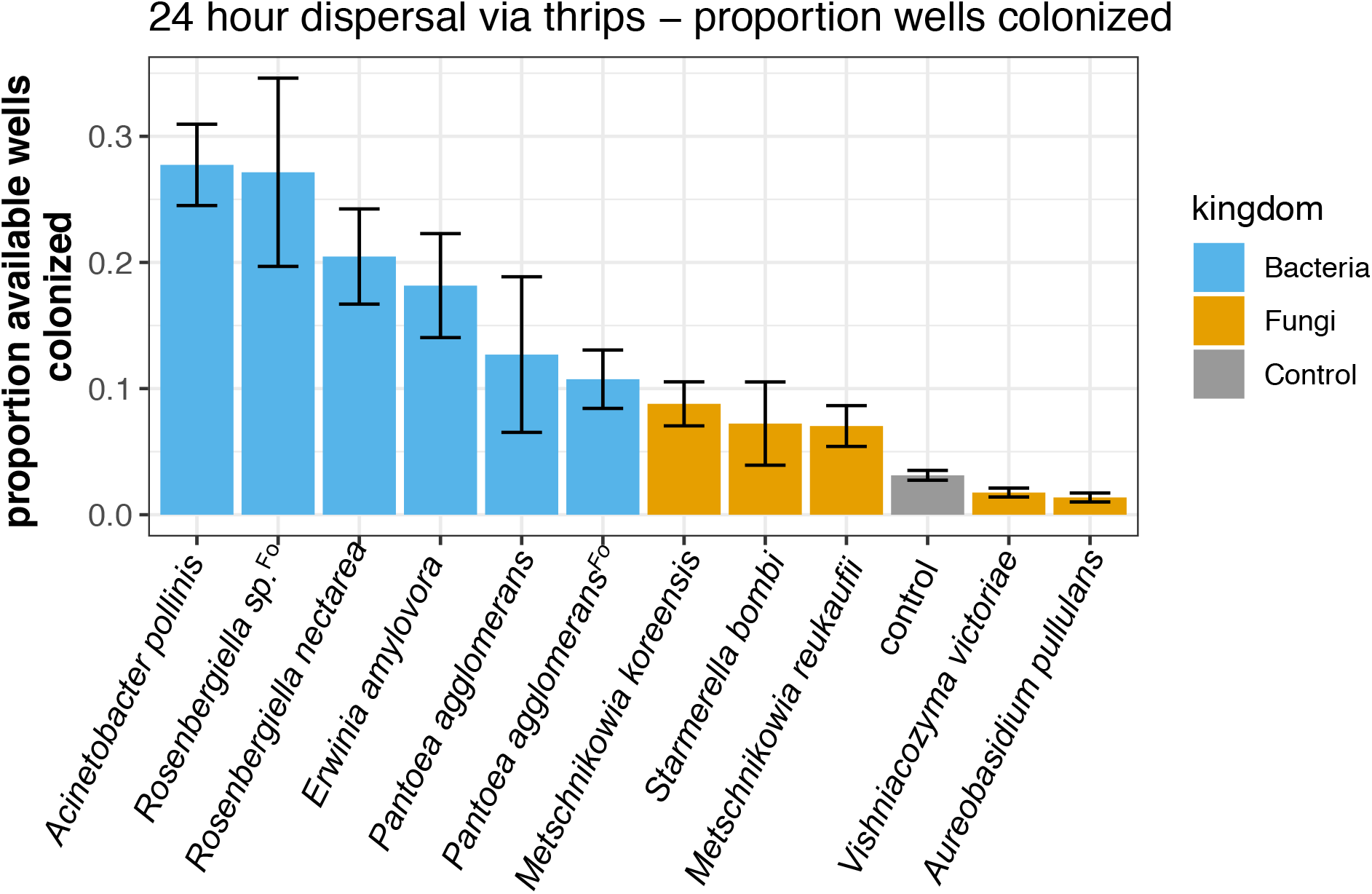
Western flower thrips (*Frankliniella occidentalis*) disperse bacteria between artificial nectar habitats more readily than fungi. Each bar shows the relative success of a different isolate of bacteria or fungus in colonizing wells on a 96-well plate through the activity of sterilized thrips (N=8-12 plates per isolate). Microbes with superscript ‘Fo’ were isolated from *Frankliniella occidentalis*. The control bar represents trials in which all wells began sterile and only thrips were added, representing contamination levels from thrips only. Bar heights report mean values (+/− 1 S.E.) for the proportion of available well’s colonized after 24 hours of thrips activity and a subsequent 6 days of culturing in an incubator.

### Patterns of bacterial and fungal incidence and co-occurrence in floral nectar

Floral nectar samples from the field contained colonies on R2A media (hereafter ‘bacteria’) in 49% of samples, whereas colonies on YMA (hereafter, ‘fungi’) were detected in only 20% of samples (microbe LRT=21.07, p<0.001, Figure 2). Fungi and bacteria co-occur in flowers nearly 1.5 times as frequently as would be expected by chance given their individual incidence across the landscape (14% of flowers contain both, compared to 9% of flowers estimated joint probability if independent, χ^2^=94.37, p<0.001). Along with the difference between bacterial (49%) and fungal (20%) incidence across the landscape, this elevated frequency of co-occurrence results in a discrepancy between the likelihoods that fungi or bacteria find themselves co-occupying a flower (conditional probabilities of co-occupancy for each microbe). Fungi (most often yeasts) are most often found in flowers alongside bacteria (P(bacteria present | fungi present) = 69%). On the other hand, bacteria only occasionally co-occupy a flower with fungi (P(fungi present | bacteria present) = 29%).

**Figure 2.**
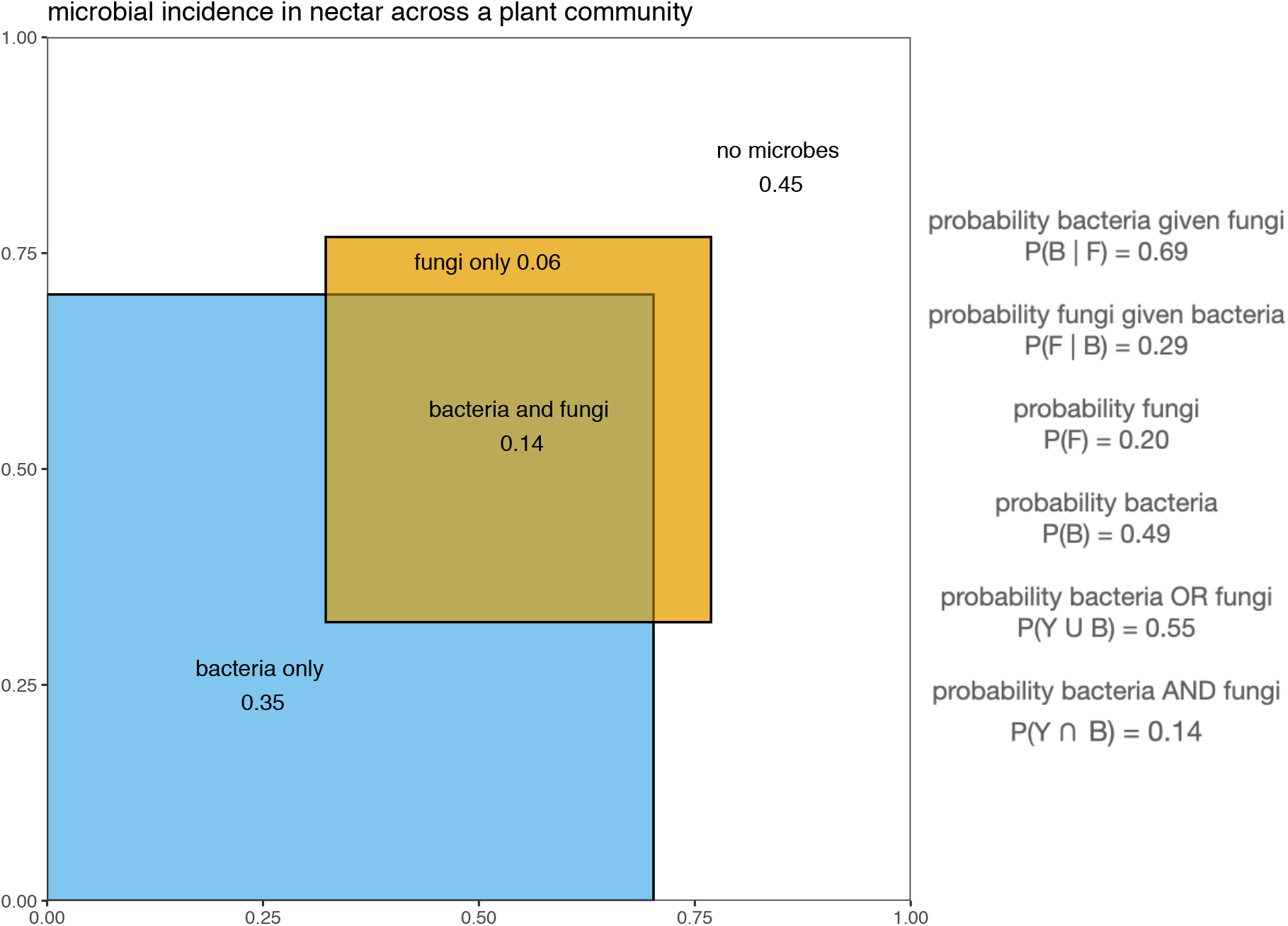
Bacterial and fungal incidence in nectar in the two flowering plant communities sampled. The figure shows the relative incidence of bacteria and fungi in individual nectar according to shaded areas and their proportions of flower occupancy. Bacteria occur in more flowers than fungi across the landscape. The two kingdoms co-occur within nectar more often than would be predicted by chance. Fungi rarely occur in the absence of bacteria, but bacteria frequently occur in the absence of fungi.

Moreover, fungi and bacteria demonstrate a positive correlation in CFU abundance across flowers (Pearson correlation, r = 0.27, p<0.001), which persists even when investigating only flowers that contain both bacteria and fungi (Pearson correlation, r=0.24, p<0.001). Within plant species, fungi and bacterial incidence and abundance are most often positively correlated, but only significantly in four species (Supplementary Figure S3).

### Nectar location associated with variation in microbial incidence and abundance

Plant species differed in microbial incidence (bacteria LRT=306, p<0.001; fungi LRT=271.09, p<0.001) and abundance (bacterial CFUs F_41,1816_=9.3, p<0.001; fungal CFUs F_41,1816_=7.3, p<0.001) in floral nectar (Figure 3) ranging from an average of 20% to 100% of sampled flowers containing bacteria. Nectar location within a flower was associated with some of this variation (Figure 4). Both bacteria and fungi were more frequently detected in nectar of plant species with an exposed nectary than those with short, mid or long corollas (Figure 4; bacteria LRT=8.1, p=0.04; fungi LRT=52.9, p<0.001). Microbial abundance also varied with flower traits, but bacteria and fungi differed in their responses. Bacteria were more abundant in flowers with exposed nectar (F_3,1813_=4.65, p=0.003) whereas fungal CFUs were most abundant when nectar is contained within a long corolla (Figure 4; F_3,1810_=29.1, p<0.001) compared to other flower morphologies.

**Figure 3.**
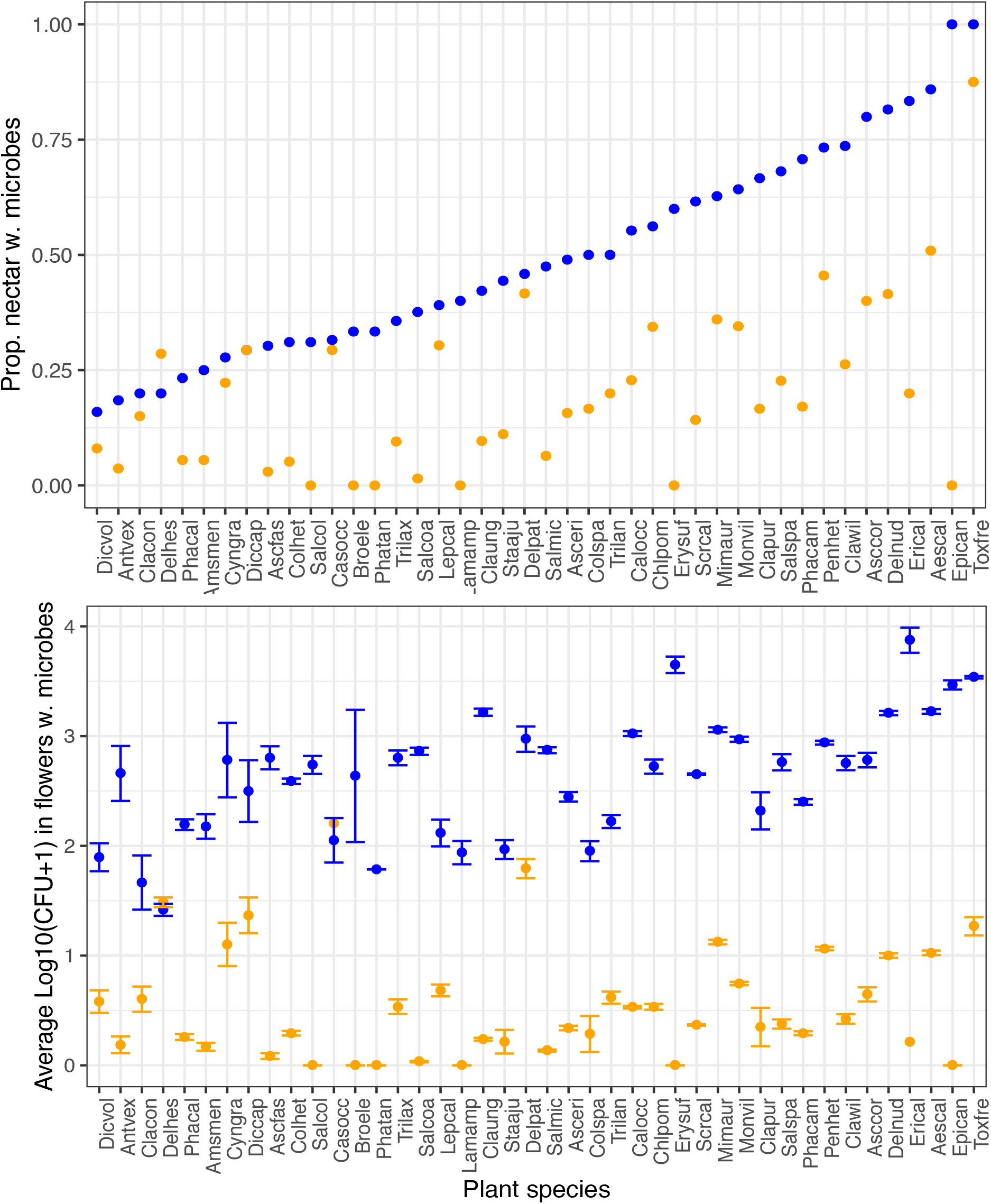
Plant species vary in A) the proportion of total nectar samples containing colonies or B) the average calculated abundance of CFUs in microbial-colonized nectar on R2A media with antifungal compound (‘bacteria’, blue) or colonies on YMA media with antibacterial (‘fungi’, orange), ordered by increasing incidence of bacteria in samples. N=6-211 per species; median=23 nectar samples/species.

**Figure 4.**
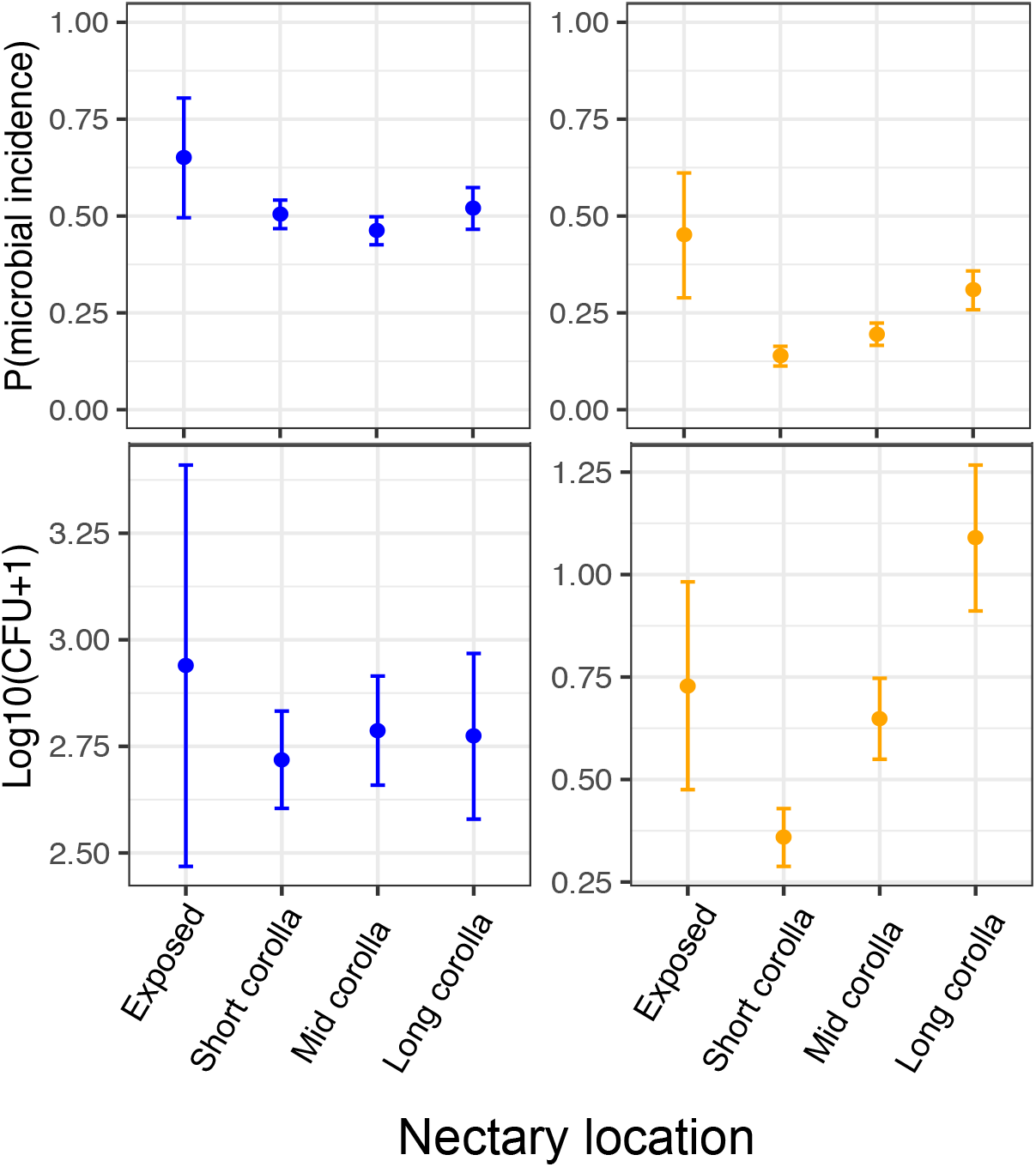
Microbial incidence and abundance in nectar varies location of the nectar (corolla length). Average microbial abundance in flowers was calculated using colony forming units counted on R2A (blue, bacteria) and YMA (fungi, orange) media and adjusted based on nectar volume. Microbial CFU abundance calculated using only those nectar samples containing detectable microbial growth. Note difference between y-axes between bacteria and fungi in microbial abundance.

### Sources of variation in bacterial and fungal incidence and abundance

Microbial incidence differed between geographic sites and years, and changed through the season (Julian date) (Supplementary Tables S3, Figure S4). Bacteria and fungi differed in which site each was more detectable and abundant, with bacteria more frequently detected and abundant at the Bee Biology site (Table S3) and fungi more frequently detected and abundant at the natural reserve site (Table S3). The probability of detection of both bacteria and fungi changed across the season—most often increasing with Julian date—and among years and sites relatively similarly (Figure S4), whereas patterns of fungal and bacterial abundance tended to differ in response to seasons and years.

### Microbial identification and genus-specific patterns

Bacteria from the genus *Acinetobacter* were the most common bacterial genera detected based on MALDI and PacBio sequencing, (Supplementary Table S4, Figure S5). Yeasts from the genera *Metschnikowia* and *Candida* were the most commonly identified fungi (Supplementary Table S4, Supplementary Figure S5). Filamentous fungi could not be identified using MALDI, but *Penicillium, Cladosporium*, and *Epicoccum* were detected using PacBio sequencing of the ITS region.

Microbial genera differed in their patterns of incidence among flowers of different morphologies. The bacterial genus *Acinetobacter* was significantly overrepresented in flowers with short corollas (Supplementary Figure S6), whereas *Pseudomonas* and the yeast *Metschnikowia* were significantly overrepresented in flowers with long corollas. Seasonal patterns were also observed: *Lactobacillus* and *Pseudomonas* were significantly overrepresented in the early season, whereas *Acinetobacter* and *Metschnikowia* were significantly underrepresented in flowers during the early season (Supplementary Figure S7).

## Discussion

In our survey, bacteria were more commonly detected in floral nectar than were fungi, and greater dispersal ability of bacteria in the experimental dispersal assay provides support for the hypothesis that differences in dispersal ability shape bacterial and fungal occupancy of coflowering communities sampled in northern California. Our use of culture-based detection is conservative in detecting bacteria (Supplementary methods S2 and Supplementary Figure S2), and the taxa detected in our sequencing efforts matched well to those detected in culture analyses, suggesting that our sampling represented well the majority of microbial community members. Our results reveal that bacteria and fungi differ in dispersal ability between flowers but frequently coexist across this and many other metacommunities. Fungal dispersal limitation may be compensated for by previously characterized differences in life histories including attraction of insect vectors by fungi or competitive exclusion of bacteria through priority effects.

Nectar-inhabiting bacteria and fungi are both likely to rely on frequent dispersal (Madden et al. 2018, Vannette 2020) because individual flowers typically persist a short time (hours to days) (Primack 1985). Previous studies and our own observations suggest multiple adaptations in both bacteria and yeasts for enhanced dispersal. Bacteria from the dominant genera found in nectar (e.g. *Acinetobacter, Rosenbergiella*)form sticky biofilms (Irie and Parsek 2008, Jung and Park 2015, Álvarez-Pérez et al. 2019), which could enhance adhesion to insect vectors. In addition, the yeasts common to nectar exhibit cell morphology that may improve adhesion to insect or bird mouthparts (e.g. ‘airplane’ cells in the yeast *Metschnikowia gruessi*(Brysch-Herzberg 2004b)). Bacteria and fungi, including yeasts, could produce stress-tolerant spores that enhance desiccation resistance (Francisco et al. 2019) and survival during transport between flowers. Microbial volatile compounds (Vannette et al. 2013, Rering et al. 2018) also affect vector attraction to flowers and visit duration (reviewed in Crowley-Gall et al. 2020). The greater dispersal ability of bacteria observed here could be due to enhanced vector attraction, ease of attachment to insect vectors, persistence in association with, survival on, or localization within hosts (Perilla-Henao and Casteel 2016), or more efficient colonization success of nectar, but further study would be required to disentangle the specific mechanisms that underlie bacterial dispersal ability.

Bacterial numerical dominance and superior dispersal ability could predict bacterial exclusion of fungi in nectar over ecological time via mass effects (Leibold et al. 2004), so it may be notable that fungi persist in the nectar environment. Coexistence between bacteria and fungi could be explained by a few factors. First, different modes of dispersal could favor different microbial species. For example, some yeasts may enhance attraction, specialize in attachment, or engage in persistent associations with particular flower visitors, as has been documented for bumble bees and *Metschnikowia* yeasts (Brysch-Herzberg 2004, Herrera et al. 2013, Jacquemyn et al. 2020). When those flower visitor species are abundant, yeast dispersal and abundance in the floral nectar metacommunity is predicted to increase. Similarly, spore-forming fungi may rely on wind dispersal, although the incidence and abundance of filamentous fungi in floral nectar was very low in our survey. Second, plant species may differ in suitability for colonization by yeasts compared to bacteria (Figures 3 & 4) or differ in flower conditions that affect yeast competitive ability. Floral nectar in some species contains antimicrobial defenses (González-Teuber and Heil 2009) including antifungal and antibacterial enzymes (Thornburg et al. 2003, Carter and Thornburg 2004, Roy et al. 2017, Schmitt et al. 2018), some of which could differentially reduce bacterial growth compared to yeast growth. Here, flowers with long corollas which host abundant yeast populations not reflected in bacterial populations. However, whether visitation by long-tongued pollinator taxa (Brysch-Herzberg 2004a, Pozo et al. 2012), or floral traits including reduced desiccation probability and reduced UV exposure (Plowright 1987) could enhance yeast colonization or growth in flowers with long corollas is unclear. Third, interference competition by fungi, including production of antimicrobial compounds, nutrient scavenging ability (Dhami et al. 2016), or environmental modification (Tucker and Fukami 2014, Vannette and Fukami 2014) could lead to suppression of bacterial populations in longer-lived flowers (but see discussion below).

Our data also reveal genus-specific patterns of plant colonization and seasonal change for a few common nectar-inhabiting microbes that may shed light on their dispersal ecology. Yeasts from the genus *Metschnikowia* were most frequently isolated from flowers with longer corollas (Supplementary Figure S6), consistent with previous reports of reliance on large-bodied bees and hummingbirds for among-flower dispersal (Herrera et al. 2010, Vannette and Fukami 2017, Morris et al. 2020). In contrast, bacteria from the genus *Acinetobacter* were more frequently isolated from flowers with shorter corollas, indicating either different dispersal vectors or the ability to successfully colonize flowers with different morphologies compared to *Metschnikowia*. For both of these nectar-inhabiting genera, incidence increased over the season. We note that many relevant ecological factors change in tandem seasonally: temperature increases, the number of flower-visiting insects increases and change in species composition, and floral density also changes across the flowering period sampled here. Previous work has shown that the flower visitor community is associated with microbial community composition and network properties in a coflowering community (Zemenick et al. 2021) but whether turnover in flower visitor species or other seasonal changes are the most important drivers of microbial composition over time is unclear.

Distinct patterns of bacterial and fungal presence and abundance in floral nectar documented here have a few specific implications for the ecology and evolution of the floral microbiome. For more prevalent nectar bacteria, inter-kingdom competition may be less important in the evolution of bacterial traits and life-history than for fungi, which are frequently exposed to competition with bacteria (Figure 2). Some models suggest that conditions of frequent yeast-bacterial interactions favor the evolution of antibacterial production by yeasts (Zhou et al. 2018). Nectar yeast have been hypothesized to produce metabolites that reduce bacterial growth (Álvarez-Pérez et al. 2019) and nectar yeast genomes contain scaffolds annotated as containing genes with potential antimicrobial function, including polyketide synthase-like and non-ribosomal peptide synthetase cluster-like (Nordberg et al. 2014), but experimental work is lacking. In contrast, experimental and genomic evidence suggests that *Metschnikowia reukaufii* benefits from rapid nutrient assimilation (Dhami et al. 2016) and population growth (Pozo et al. 2012), suggesting maximizing propagule abundance is an important strategy for nectar microbial populations (Hibbing et al. 2010). Moreover, our data do not reveal strong negative correlations between fungi and bacterial abundance in nectar, either among or within species (Supplementary Figure S3), suggesting that interference competition may not be strong in our focal populations. Instead, we noted positive correlations between fungal and bacterial abundance at the flower level, which could indicate shared dispersal vectors, similar habitat requirements for culturable nectar microbes, or the possibility of facilitation which could be promoted by bacterial enhancement of nutrients in nectar from pollen or other plant sources (Christensen et al. 2021).

Taken together, the data presented here demonstrate that common nectar-inhabiting fungi are more dispersal limited than are bacteria and less frequently detected among flowers. Previous work has shown that dispersal can be important in structuring bacterial community assembly (Albright and Martiny 2018), and generate the possibility of historically contingent competitive outcomes in microbial communities (Fukami 2015, Svoboda et al. 2018). By demonstrating clear differences in dispersal ability, our results extend this to interactions between bacteria and fungi. We suggest that further study of the consequences of variation in dispersal differences between bacterial and fungal taxa may inform the types and temporal dynamics of interactions between them, and consequences for host-microbe and multitrophic interactions in flowers.

## Author contributions

RLV conceived of the field study, performed fieldwork, and wrote the manuscript. MM and RLV conceived of the thrips dispersal assay and performed statistical analyses. GH performed field and microbial labwork. MM and DP performed the thrips dispersal assay. IM analyzed and processed samples for MALDI-TOF identifications. TM performed dilution plating assays. All authors contributed to revisions and gave final approval for publication.

## Acknowledgements

We are grateful to the University of California Natural Reserve System, the Laidlaw Honey Bee Research Facility, Jeffrey Clarey and Neal Williams for site access, Diane Ullman for the thrips colony, Amin Montazer for help with lab work, Will Jewell and Barbara Byrnes for training and technical support with MALDI-TOF, and to the Vannette lab group for providing feedback on initial drafts of this manuscript. RLV was supported by UC Davis, the Hellman Fund, the United States Department of Agriculture Hatch funds multistate NE1501 and the National Science Foundation DEB #1846266.

## Data and code accessibility statement

Data and code are available through Dryad DOI: https://doi.org/10.25338/B8403V

## References

Albright, M. B. N., and J. B. H. Martiny. 2018. Dispersal alters bacterial diversity and composition in a natural community. The ISME journal 12:296–299.

Alexander, H. M., and J. Antonovics. 1988. Disease spread and population dynamics of anther-smut infection of Silene alba caused by the fungus Ustilago violacea. The Journal of Ecology:91–104.

Alvarez-Pérez, S., and C. M. Herrera. 2013. Composition, richness and nonrandom assembly of culturable bacterial-microfungal communities in floral nectar of Mediterranean plants. FEMS Microbiol Ecol 83.

Álvarez-Pérez, S., C. M. Herrera, and C. de Vega. 2012. Zooming-in on floral nectar: a first exploration of nectar-associated bacteria in wild plant communities. Fems Microbiology Ecology 80:591–602.

Álvarez-Pérez, S., B. Lievens, and T. Fukami. 2019. Yeast–Bacterium Interactions: The Next Frontier in Nectar Research. Trends in plant science 24:393–401.

Baguette, M., T. G. Benton, and J. M. Bullock. 2012. Dispersal ecology and evolution. Oxford University Press.

Barberán, A., J. Ladau, J. W. Leff, K. S. Pollard, H. L. Menninger, R. R. Dunn, and N. Fierer. 2015. Continental-scale distributions of dust-associated bacteria and fungi. Proceedings of the National Academy of Sciences 112:5756–5761.

Belisle, M., K. G. Peay, and T. Fukami. 2012. Flowers as islands: spatial distribution of nectar-inhabiting microfungi among plants of *Mimulus aurantiacus*, a hummingbird-pollinated shrub. Microbial Ecology 63:711–718.

Brysch-Herzberg, M. 2004a. Ecology and taxonomy of yeasts associated with the plant-bumblebee mutualism in central Europe. FEMS Microbiol Ecol 50.

Brysch-Herzberg, M. 2004b. Ecology of yeasts in plant-bumblebee mutualism in Central Europe. Fems Microbiology Ecology 50:87–100.

Callahan, B. J., P. J. McMurdie, M. J. Rosen, A. W. Han, A. J. A. Johnson, and S. P. Holmes. 2016. DADA2: high-resolution sample inference from Illumina amplicon data. Nature methods 13:581.

Callahan, B. J., J. Wong, C. Heiner, S. Oh, C. M. Theriot, A. S. Gulati, S. K. McGill, and M. K. Dougherty. 2019. High-throughput amplicon sequencing of the full-length 16S rRNA gene with single-nucleotide resolution. Nucleic acids research 47:e103–e103.

Carter, C., and R. W. Thornburg. 2004. Is the nectar redox cycle a floral defense against microbial attack? Trends in Plant Science 9:320–324.

Chappell, C. R., and T. Fukami. 2018. Nectar yeasts: a natural microcosm for ecology. Yeast 35:417–423.

Christensen, S. M., I. Munkres, and R. L. Vannette. 2021. Nectar bacteria stimulate pollen germination and bursting to enhance their fitness. bioRxiv:2021.2001.2007.425766.

Crowley-Gall, A., C. R. Rering, A. B. Rudolph, R. L. Vannette, and J. J. Beck. 2020. Volatile microbial semiochemicals and insect perception at flowers. Current Opinion in Insect Science.

Dhami, M. K., T. Hartwig, and T. Fukami. 2016a. Genetic basis of priority effects: insights from nectar yeast. Proceedings of the Royal Society B: Biological Sciences 283:20161455.

Dhami, M. K., T. Hartwig, A. D. Letten, M. Banf, and T. Fukami. 2018. Genomic diversity of a nectar yeast clusters into metabolically, but not geographically, distinct lineages. Molecular Ecology 27:2067–2076.

Francisco, C. S., X. Ma, M. M. Zwyssig, B. A. McDonald, and J. Palma-Guerrero. 2019. Morphological changes in response to environmental stresses in the fungal plant pathogen Zymoseptoria tritici. Scientific reports 9:9642.

Fridman, S., I. Izhaki, Y. Gerchman, and M. Halpern. 2012. Bacterial communities in floral nectar. Environmental Microbiology Reports 4:97–104.

Fukami, T. 2015. Historical contingency in community assembly: integrating niches, species pools, and priority effects. Annual Review of Ecology, Evolution, and Systematics 46.

González-Teuber, M., and M. Heil. 2009. Nectar chemistry is tailored for both attraction of mutualists and protection from exploiters. Plant signaling & behavior 4:809–813.

Herrera, C. M., A. Canto, M. I. Pozo, and P. Bazaga. 2010. Inhospitable sweetness: nectar filtering of pollinator-borne inocula leads to impoverished, phylogenetically clustered yeast communities. Proceedings of the Royal Society B: Biological Sciences 277:747–754.

Herrera, C. M., C. de Vega, A. Canto, and M. I. Pozo. 2009. Yeasts in floral nectar: a quantitative survey. Annals of Botany 103:1415–1423.

Herrera, C. M., I. M. Garcia, and R. Perez. 2008. Invisible floral larcenies: Microbial communities degrade floral nectar of bumble bee-pollinated plants. Ecology 89:2369–2376.

Herrera, C. M., M. I. Pozo, and M. Medrano. 2013. Yeasts in nectar of an early-blooming herb: sought by bumble bees, detrimental to plant fecundity. Ecology 94:273–279.

Hibbing, M. E., C. Fuqua, M. R. Parsek, and S. B. Peterson. 2010. Bacterial competition: surviving and thriving in the microbial jungle. Nature reviews. Microbiology 8:15–25.

Irie, Y., and M. R. Parsek. 2008. Quorum Sensing and Microbial Biofilms. Pages 67–84 in T. Romeo, editor. Bacterial Biofilms. Springer Berlin Heidelberg, Berlin, Heidelberg.

Jacquemyn, H., M. I. Pozo, S. Álvarez-Pérez, B. Lievens, and T. Fukami. 2020. Yeast-nectar interactions: metacommunities and effects on pollinators. Current Opinion in Insect Science.

Johnson, R. M., J. D. Evans, G. E. Robinson, and M. R. Berenbaum. 2009. Changes in transcript abundance relating to colony collapse disorder in honey bees *(Apis mellifera)*. Proceedings of the National Academy of Sciences 106:14790–14795.

Jung, J., and W. Park. 2015. Acinetobacter species as model microorganisms in environmental microbiology: current state and perspectives. Applied Microbiology and Biotechnology 99:2533–2548.

Kingston, A. B., and P. B. Mc Quillan. 2000. Are pollination syndromes useful predictors of floral visitors in Tasmania? Austral Ecology 25:600–609.

Kõljalg, U., K. H. Larsson, K. Abarenkov, R. H. Nilsson, I. J. Alexander, U. Eberhardt, S. Erland, K. Høiland, R. Kjøller, and E. Larsson. 2005. UNITE: a database providing web-based methods for the molecular identification of ectomycorrhizal fungi. New Phytologist 166:1063–1068.

Koskella, B., L. J. Hall, and C. J. E. Metcalf. 2017. The microbiome beyond the horizon of ecological and evolutionary theory. Nature Ecology & Evolution 1:1606–1615.

Kraemer, S., and P. Boynton. 2016. Evidence for microbial local adaptation in nature. Molecular Ecology.

Lachance, M. A., W. T. Starmer, C. A. Rosa, J. M. Bowles, J. S. F. Barker, and D. H. Janzen. 2001. Biogeography of the yeasts of ephemeral flowers and their insects. Fems Yeast Research 1:1–8.

Lara, C., and J. Ornelas. 2001. Preferential nectar robbing of flowers with long corollas: experimental studies of two hummingbird species visiting three plant species. Oecologia 128:263–273.

Leibold, M. A., M. Holyoak, N. Mouquet, P. Amarasekare, J. Chase, M. Hoopes, R. Holt, J. Shurin, R. Law, and D. Tilman. 2004. The metacommunity concept: a framework for multi-scale community ecology. Ecology Letters 7:601–613.

Lenaerts, M., T. Goelen, C. Paulussen, B. Herrera-Malaver, J. Steensels, W. Van den Ende, K. J. Verstrepen, F. Wäckers, H. Jacquemyn, and B. Lievens. 2017. Nectar bacteria affect life history of a generalist aphid parasitoid by altering nectar chemistry. Functional Ecology.

Lindow, S. E., and G. L. Andersen. 1996. Influence of immigration on epiphytic bacterial populations on navel orange leaves. Applied and environmental microbiology 62:2978–2987.

Madden, A. A., M. J. Epps, T. Fukami, R. E. Irwin, J. Sheppard, D. M. Sorger, and R. R. Dunn. 2018. The ecology of insect-yeast relationships and its relevance to human industry. Proceedings of the Royal Society B: Biological Sciences 285:2017–2733.

Mittelbach, M., A. M. Yurkov, R. Stoll, and D. Begerow. 2016. Inoculation order of nectar-borne yeasts opens a door for transient species and changes nectar rewarded to pollinators. Fungal Ecology.

Morris, M., N. Frixione, A. Burkert, E. Dinsdale, and R. L. Vannette. 2020. Microbial abundance, composition, and function in nectar are shaped by flower visitor identity. FEMS Microbiol Ecol 96:3.

Mouquet, N., and M. Loreau. 2003. Community patterns in source-sink metacommunities. The american naturalist 162:544–557.

Ngugi, H. K., and H. Scherm. 2006. Biology of flower-infecting fungi. Annu. Rev. Phytopathol. 44:261–282.

Nordberg, H., M. Cantor, S. Dusheyko, S. Hua, A. Poliakov, I. Shabalov, T. Smirnova, I. V. Grigoriev, and I. Dubchak. 2014. The genome portal of the Department of Energy Joint Genome Institute: 2014 updates. Nucleic acids research 42:D26–D31.

Peay, K. G., M. G. Schubert, N. H. Nguyen, and T. D. Bruns. 2012. Measuring ectomycorrhizal fungal dispersal: macroecological patterns driven by microscopic propagules. Molecular Ecology 21:4122–4136.

Perilla-Henao, L. M., and C. L. Casteel. 2016. Vector-Borne Bacterial Plant Pathogens: Interactions with Hemipteran Insects and Plants. Frontiers in Plant Science 7.

Plowright, R. 1987. Corolla depth and nectar concentration: an experimental study. Canadian Journal of Botany 65:1011–1013.

Pozo, M. I., M.-A. Lachance, and C. M. Herrera. 2012. Nectar yeasts of two southern Spanish plants: the roles of immigration and physiological traits in community assembly. Fems Microbiology Ecology 80:281–293.

Primack, R. B. 1985. Longevity of individual flowers. Annual Review of Ecology and Systematics 16:15–37.

Quast, C., E. Pruesse, P. Yilmaz, J. Gerken, T. Schweer, P. Yarza, J. Peplies, and F. O. Glöckner. 2012. The SILVA ribosomal RNA gene database project: improved data processing and web-based tools. Nucleic acids research 41:D590–D596.

Rering, C. C., J. J. Beck, G. W. Hall, M. M. McCartney, and R. L. Vannette. 2018. Nectar-inhabiting microorganisms influence nectar volatile composition and attractiveness to a generalist pollinator. New Phytologist.

Roy, R., A. J. Schmitt, J. B. Thomas, and C. J. Carter. 2017. Nectar biology: from molecules to ecosystems. Plant Science 262:148–164.

Schaeffer, R., and R. E. Irwin. 2014. Yeasts in nectar enhance male fitness in a montane perennial herb. Ecology 95:1792–1798.

Schmitt, A. J., A. E. Sathoff, C. Holl, B. Bauer, D. A. Samac, and C. J. Carter. 2018. The major nectar protein of Brassica rapa is a non-specific lipid transfer protein, BrLTP2.1, with strong antifungal activity. Journal of Experimental Botany 69:5587–5597.

Sobhy, I. S., D. Baets, T. Goelen, B. Herrera-Malaver, L. Bosmans, W. Van den Ende, K. J. Verstrepen, F. Wäckers, H. Jacquemyn, and B. Lievens. 2018. Sweet Scents: Nectar Specialist Yeasts Enhance Nectar Attraction of a Generalist Aphid Parasitoid Without Affecting Survival. Frontiers in Plant Science 9.

Svoboda, P., E. S. Lindström, O. Ahmed Osman, and S. Langenheder. 2018. Dispersal timing determines the importance of priority effects in bacterial communities. The ISME journal 12:644–646.

Thornburg, R. W., C. Carter, A. Powell, R. Mittler, L. Rizhsky, and H. T. Horner. 2003. A major function of the tobacco floral nectary is defense against microbial attack. Plant Systematics and Evolution 238:211–218.

Toju, H., R. L. Vannette, M. P. L. Gauthier, M. K. Dhami, and T. Fukami. 2018. Priority effects can persist across floral generations in nectar microbial metacommunities. Oikos 127:345–352.

Tsuji, K., and T. Fukami. 2018. Community-wide consequences of sexual dimorphism: evidence from nectar microbes in dioecious plants. Ecology 99:2476–2484.

Tucker, C. M., and T. Fukami. 2014. Environmental variability counteracts priority effects to facilitate species coexistence: evidence from nectar microbes. Proceedings of the Royal Society of London B: Biological Sciences 281:2013–2637.

Vannette, R. L. 2020. The Floral Microbiome: Plant, Pollinator, and Microbial Perspectives. Annual Review of Ecology, Evolution, and Systematics 51:363–386.

Vannette, R. L., and T. Fukami. 2014. Historical contingency in species interactions: towards niche-based predictions. Ecology letters 17:115–124.

Vannette, R. L., and T. Fukami. 2017. Dispersal enhances beta diversity in nectar microbes. Ecology Letters.

Vannette, R. L., M.-P. L. Gauthier, and T. Fukami. 2013. Nectar bacteria, but not yeast, weaken a plant-pollinator mutualism. Proceedings of the Royal Society B-Biological Sciences 280.

von Arx, M., A. Moore, G. Davidowitz, and A. E. Arnold. 2019. Diversity and distribution of microbial communities in floral nectar of two night-blooming plants of the Sonoran Desert. Plos One 14.

Zhou, N., M. Katz, W. Knecht, C. Compagno, and J. Piškur. 2018. Genome dynamics and evolution in yeasts: A long-term yeast-bacteria competition experiment. Plos One 13:e0194911.

