## Supplementary materials for "Fungi are more dispersal limited than bacteria among flowers"

Supplementary Material for “Fungi are more dispersal-limited than bacteria among flowers”

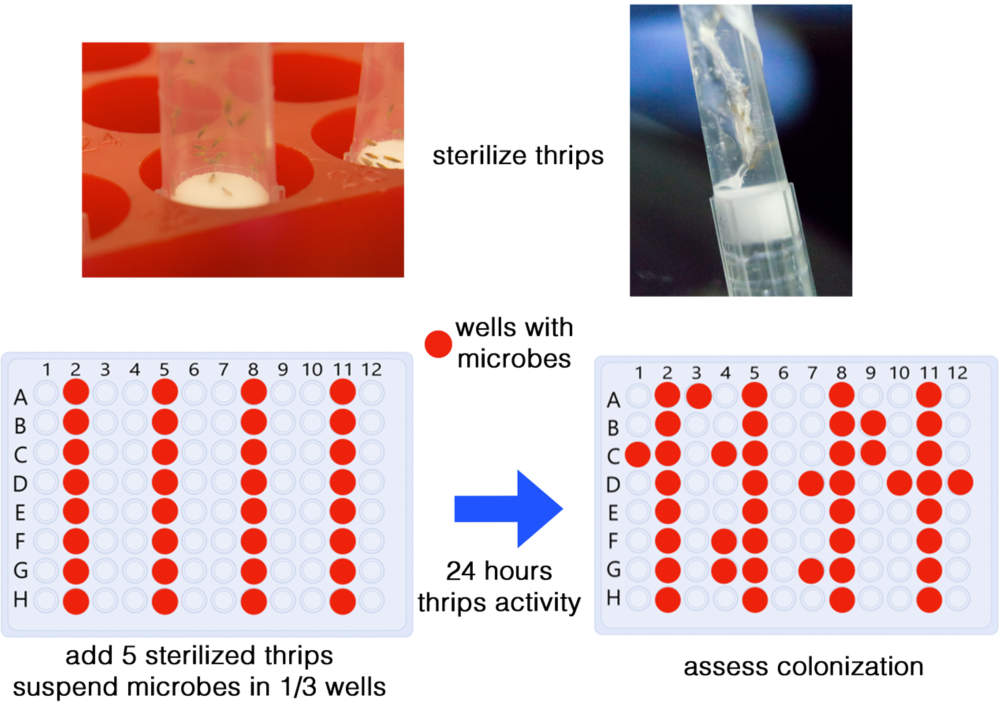


**Supplemental Figure S1**

Thrips (*Frankliniella occidentalis)* were first fed a diet containing the antibacterial chloramphenicol. Thrips were sterilized in batches through repeated washings inside pipette chambers. Washing consisted of vortexing at maximum rpm and plunging out solution with ethanol, water, bleach, and water three more times. The assay layout used included 1/3 of wells beginning with microbes and waiting 24 hours for thrips to disperse microbes.

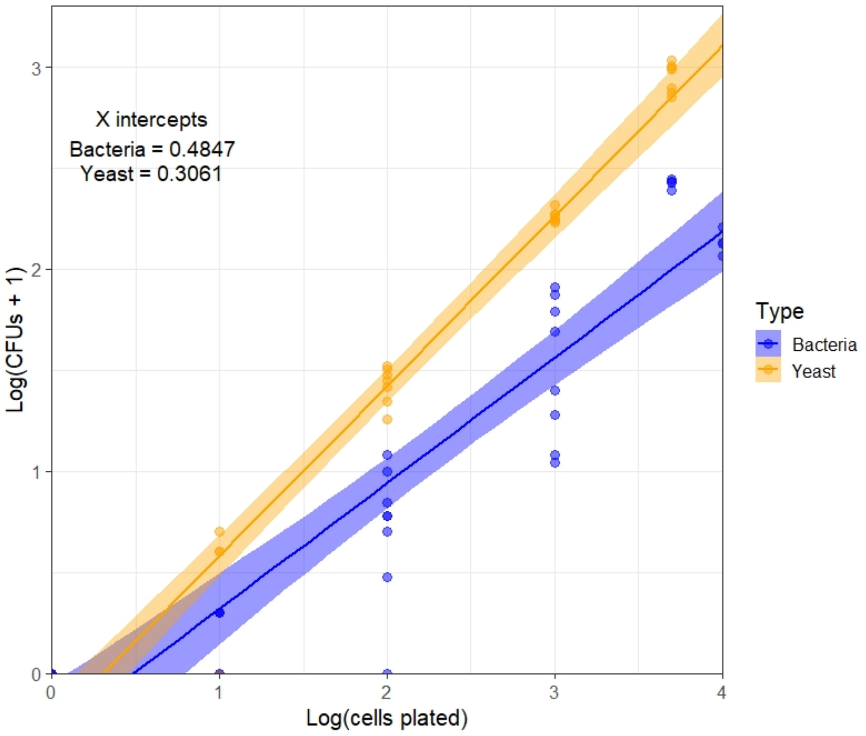


**Supplementary Figure S2**—Relationship between cell number in suspension (plated) and CFU detection on a) Reasoner's Agar (R2A) and b) and yeast agar media (YMA) plates, using the bacteria *Acinetobacter nectaris* and the yeast *Metschnikowia reukaufii* for R2A and YMA plates, respectively.

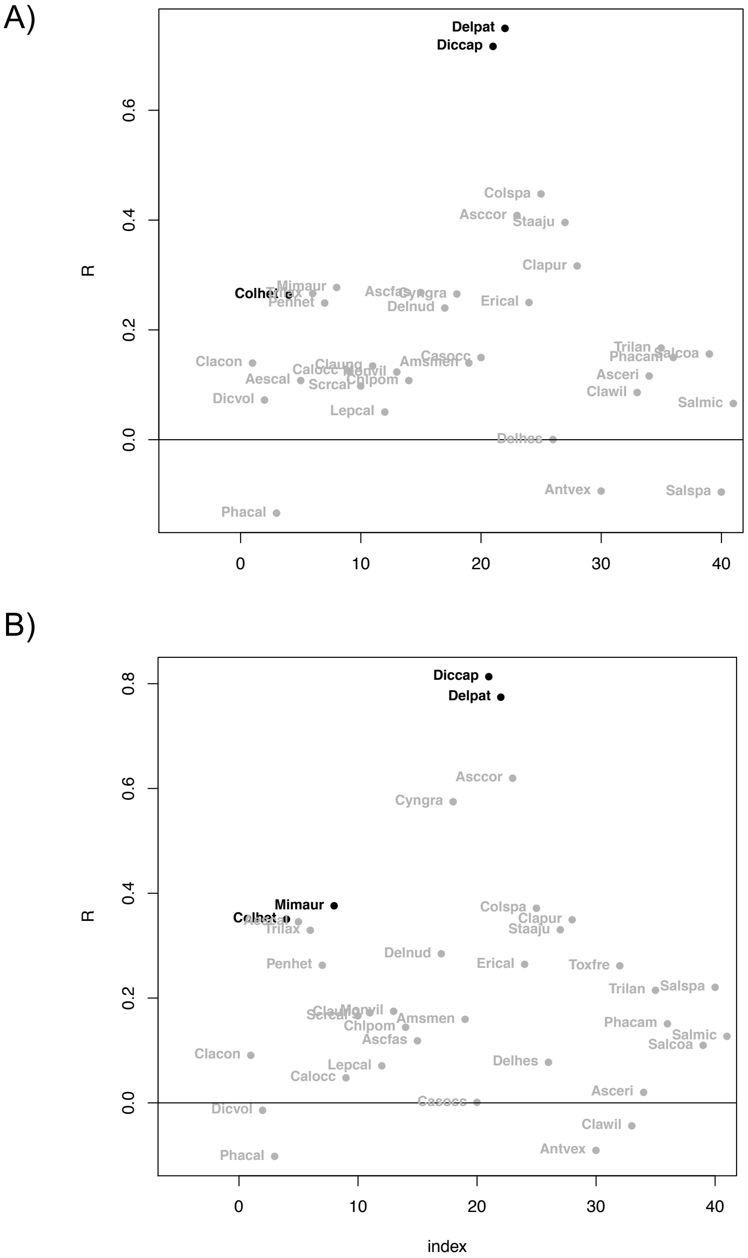


**Supplementary Figure S3.** Within-species correlation coefficients for Pearson correlations between bacterial and fungal A) presence absence and B) abundance. Points are colored black when the correlation is significant at FDR >0.05 and labeled with genus and species abbreviations. Note that all but four correlation coefficients are positive (above 0).

A)

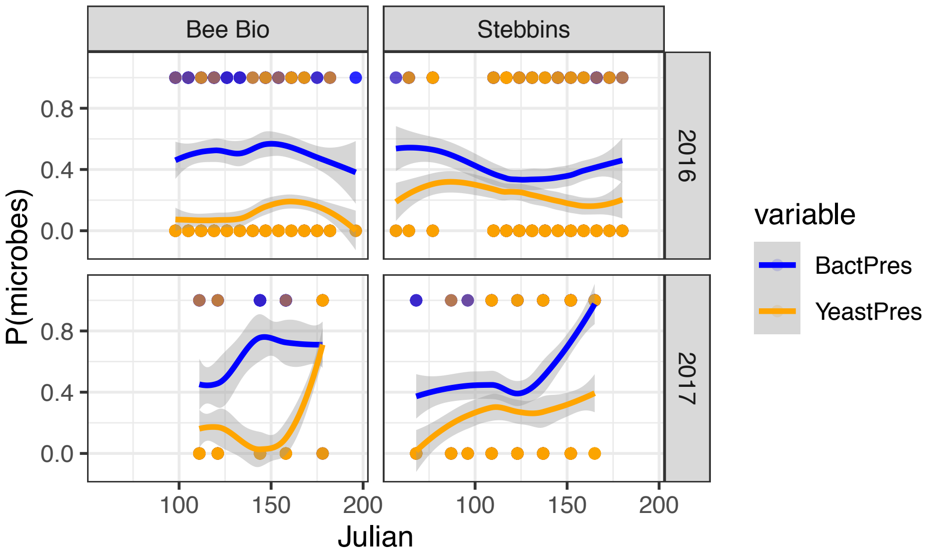


B)

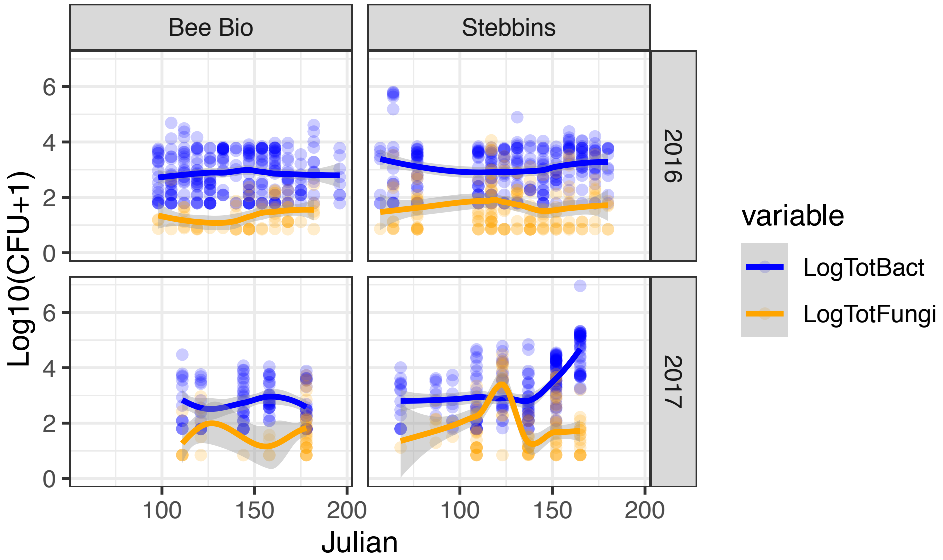


**Supplementary Figure S4.** Seasonal patterns in the a) incidence of and b) abundance of colony forming units on Reasoner's agar (R2A) plates (bacteria, blue) and colonies on yeast media agar plates (YMA, fungi, in orange) found in floral nectar from two sampling sites in Northern California. In panel b) points are only shown for non-zero values. Sites were sampled through the duration of the main spring flowering period at both sites during 2016 and 2017. Lines represent loess fit with 95% confidence values. Points represent individual nectar samples.

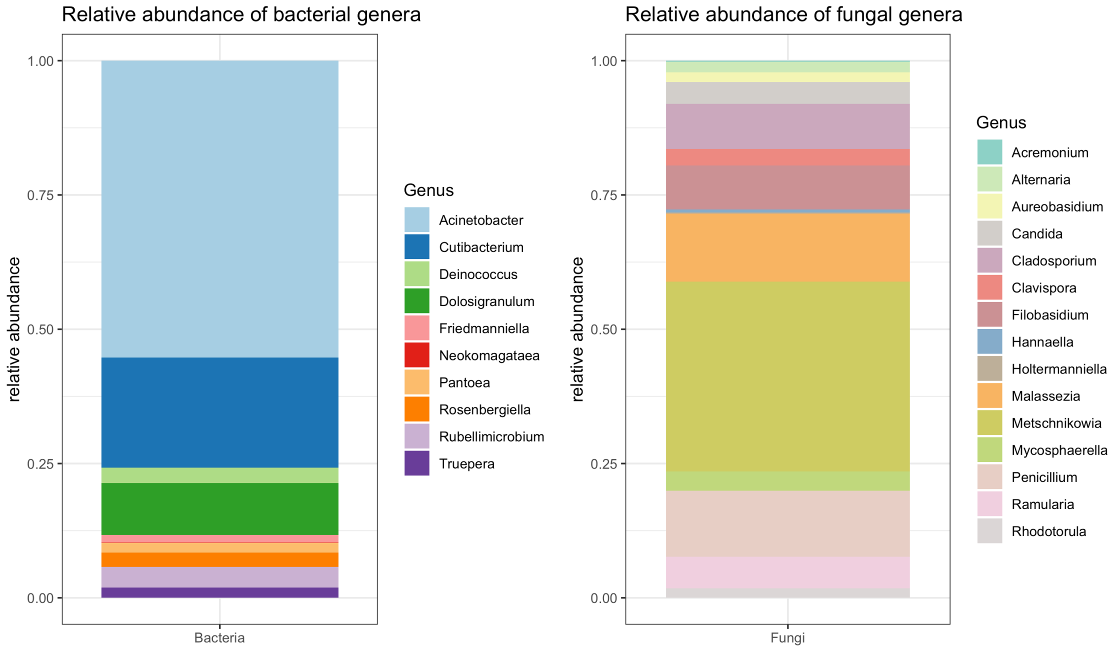


**Supplementary Figure S5**. Summary of taxa detected in pooled nectar samples. Bacterial data were recovered from N=7 samples for 16S and N=17 samples for ITS sequences. Chloroplast contamination accounted for 37% of reads in the ITS reads and 30% of reads in the 16S dataset.

**Supplementary Figure S6.** Frequency of detection for most abundant microbial genera found in floral nectar and identified by MALDI, plotted by flower morphology of plant species from which it was isolated. Blue bars indicate that taxa was more frequently detected than expected by chance and red bars indicate that taxa was less frequently detected than expected by chance. Bar thickness is proportional to the number of times this species was detected in each flower types. P-value indicates significance based on an independence model for a contingency table.

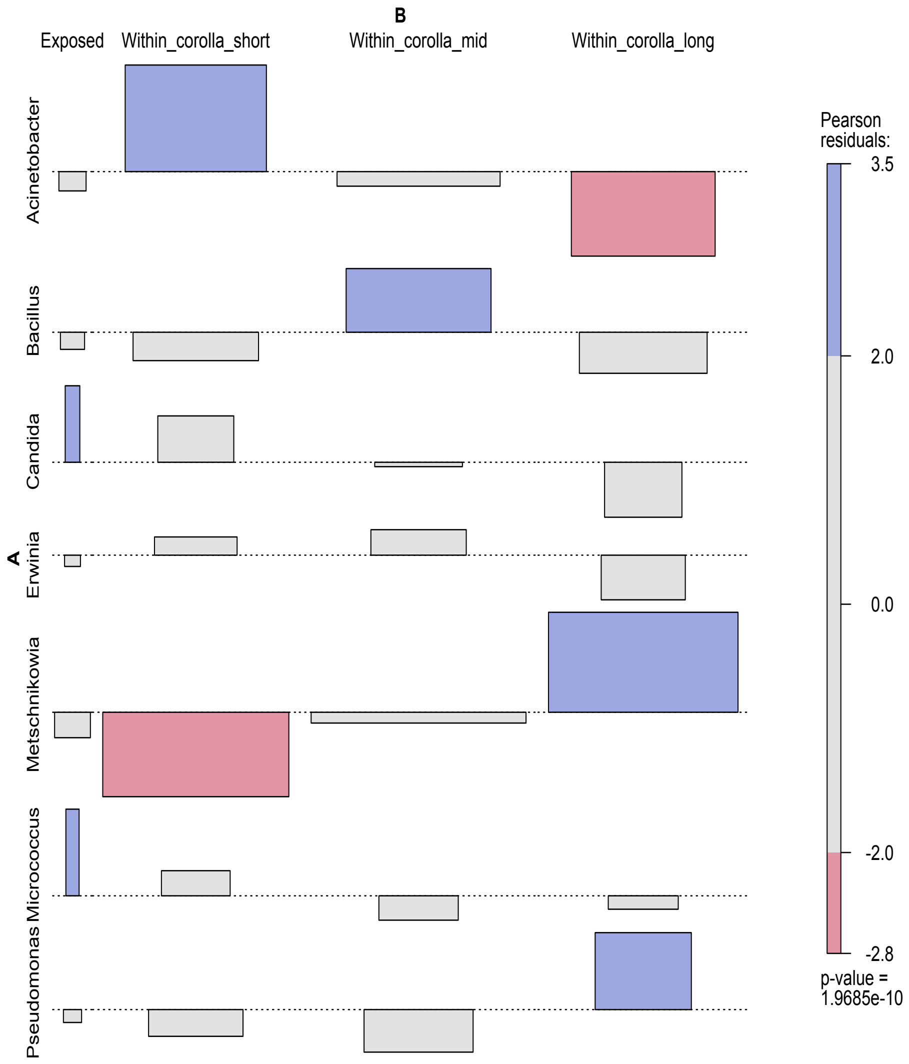

**Supplementary Figure S7**. Frequency of detection for most abundant microbial genera found in floral nectar and identified by MALDI, plotted by time of the flowering season, defined by Julian date, where ‘early’<100, 100≤‘mid’<150, and ‘late’≥150. Blue bars indicate that taxa was more frequently detected than expected by chance and red bars indicate that taxa was less frequently detected than expected by chance. Bar thickness is proportional to the number of times this species was detected. P-value indicates significance based on an independence model for a contingency table.

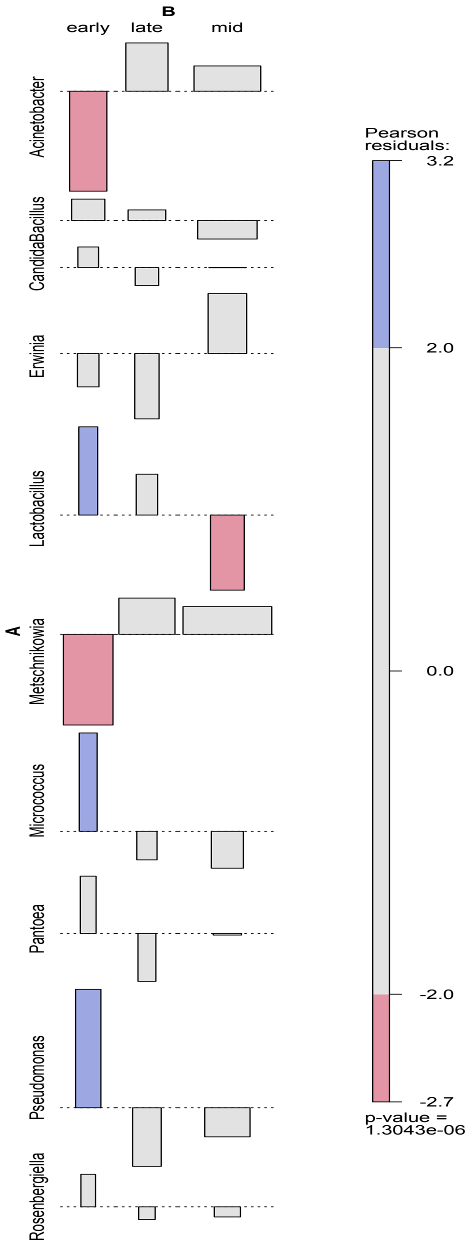

**Supplementary Table S1.** Microbial strains used in dispersal assays, closest BLAST match for species and genus, host species sources, and identifying sequences. Primer sequences used to amplify yeasts were NL1/NL4 or ITS1f/ITS4; bacteria were amplified using 27F/1492R.

| *Strain ID* | *Kingdom* | *species_Genus* | *Host plant or insect source* | *sequence* |
|---|---|---|---|---|
| *EC52* | F | Metschnikowia_reukaufii | Epilobium_canum | CCTTCGGGAATTGTATTTTGAAGGTGGGTTTGGTTAGGAAAAGTTACTTTAAGTCCATTGGAAAATGGCGCCATGGAGGGTGATAGCCCCGTAAAAGTATCCCTTTTCCTTTTATCCATTCCCTCCAAAGAGTCGAGTTGTTTGGGAATGCAGCTCTAAGTGGGTGGTAAATTCCATCTAAAGCTAAATATTGGCGAGAGACCGATAGCGAACAAGTACAGTGATGGAAAGATGAAAAGCACTTTGAAAAGAGAGTGAAAAAGTACGTGAAATTGTTGAAAGGGAAGGGCTTGCAAGCAGACACAACCTCGGTTGGGCCAGCATCGGAGTGGGGGGAGACAAAAAAGAAAAGGAATGTAACTCATTGAGTATTATAGCCTTTTTCTCATATCTCCACCCCCTTCCG |
| *EC124* | B | Rosenbergiella_nectarea | Epilobium_canum | TCTTTTGCAACCCACTCCCATGGTGTGACGGGCGGTGTGTACAAGGCCCGGGAACGTATTCACCGTAACATTCTGATTTACGATTACTAGCGATTCCGACTTCATGGAGTCGAGTTGCAGACTCCAATCCGGACTACGACGCACTTTATGAGGTCCGCTTGCTCTCGCGAGGTCGCTTCTCTTTGTATGCGCCATTGTAGCACGTGTGTAGCCCTACTCGTAAGGGCCATGATGACTTGACGTCATCCCCACCTTCCTCCGGTTTATCACCGGCAGTCTCCTTTGAGTTCCCACCATTACGTGCTGGCAACAAAGGATAAGGGTTGCGCTCGTTGCGGGACTTAACCCAACATTTCACAACACGAGCTGACGACAGCCATGCAGCACCTGTCTCAGAGTTCCCGAAGGCACTAAAGCATCTCTGCTAAATTCTCTGGATGTCAAGAGTAGGTAAGGTTCTTCGCGTTGCATCGAATTAAACCACATGCTCCACCGCTTGTGCGGGCCCCCGTCAATTCATTTGAGTTTTAACCTTGCGGCCGTACTCCCCA |
| *FO-03* | B | Rosenbergiella_sp | Frankliniella_occidentalis | CTTGCTACTTTGCTGACGAGTGGCGGACGGGTGAGTAATGTCTGGGGATCTGCCTGATGGAGGGGGATAACTACTGGAAACGGTAGCTAATACCGCATAATGTCGCAAGACCAAAGCGGGGGACTTTCGGGCCTCGCACCATCAGATGAACCCAGATGGGATTAGCTAGTAGGTAAGGTAATGGCTTACCTAGGCGACGATCCCTAGCTGGTCTGAGAGGATGACCAGCCACACTGGAACTGAGACACGGTCCAGACTCCTACGGGAGGCAGCAGTGGGGAATATTGCACAATGGGCGCAAGCCTGATGCAGCCATGCCGCGTGTATGAAGAAGGCCTTCGGGTTGTAAAGTACTTTCAGTCAGGAGGAAGGGTGTGAAATTAATACTTTCATGCATTGACGTTACTGACAGAAGAAGCACCGGCTAACTCCGTGCCAKC |
| *SCC477* | B | Acinetobacter_pollinis | Scrophularia_californica | TACTAGCGATTCCGACTTCATGGAGTCGAGTTGCAGACTCCAATCCGGACTACGATCGGCTTTTTGAGATTAGCATCACATCGCTGTGTAGCAACCCTCTGTACCGACCATTGTAGCACGTGTGTAGCCCTGGCCGTAAGGGCCATGATGACTTGACGTCGTCCCCGCCTTCCTCCAGTTTGTCACTGGCAGTATCCTTAAAGTTCCCATCCGAAATGCTGGCAAGTAAGGAAAAGGGTTGCGCTCGTTGCGGGACTTAACCCAACATCTCACGACACGAGCTGACGACAGCCATGCAGCACCTGTATCTAAGTTCCCGAAGGCACCAATCTATCTCTAGAAAGTTCTTAGTATGTCAAGGCCAGGTAAGGTTCTTCGCGTTGCATCGAATTAAACCACATGCTCCACCGCTTGTGCGGGCCCC |

| | | | | |
|---|---|---|---|---|
| | | | | CGTCAATTCATTTGAGTTTTAGTCTTGCGACCGTACTCCCCAGGCGGTCTACTTATCGCGTTAGCTGCGCCACTAAGTCCTCAAAGGACCCAACGGCT |
| *FO-01* | B | Pantoea_agglomerans | Frankliniella_occidentalis | TCTTTTGCAACCCACTCCCATGGTGTGACGGGCGGTGTGTACAAGGCCCGGGAACGTATTCACCGTGGCATTCTGAKCCAYGATYAMAMTCGYWTCCGACTTCACGGAGTCGAGTTGCAGACTCCGATCCGGACTACGACGCACTTTGTGAGGTCCGCTTGCTCTCGCGAGGTCGCTTCTCTTTGTATGCGCCATTGTAGCACGTGTGTAGCCCTACTCGTAAGGGCCATGATGACTTGACGTCATCCCCACCTTCCTCCGGTTTATCACCGGCAGTCTCCTTTGAGTTCCCGACCGAATCGCTGGCAACAAAGGATAAGGGTTGCGCTCGTTGCGGGACTTAACCCAACATTTCACAACACGAGCTGACGACAGCCATGCAGCACCTGTCTCASSGTTCCCGAAGGCACYAAAGCATCTCTGC |
| *Q1F2* | F | Starmerella_bombi | Bombus_vosnesenskii | AACTCGCATTCGGAGAGTCATGCCGTACTCGTCGTCCGCTTAGTGTGATATAGCGACTGAAGCTATAATACTCCGAGGAGTTACATTCTTCAGCTTTTATCTTCCCCCCAAACACGACTCTACGTGGTTAGCGGCCTACCCTTCCATTTCAACAATTTCACGTACTTTTTCACTCTCTTTTCAAAGTTCTTTTCATCTTTCCTTCACAGTACTTGTTCGCTATCGGTCTCTCGCAGATATTTAGCTTTAGATGGAGCATACCACCCATTTGAGCTGCATTCCCAAACAACTCGACTCCATGCCAAGGTCCTACAGTGGGGTCAATGTCGTACGGGGCTATCACCCTCCATGGCGCTCCTTTCCAGAAGACTTAGACATCGGTTTCCCAGGACCAAGGCTTCAGAATACAATGCCCCGAAAGGCTTTCAAATCTGAGCTCTTGCCTGTTCACTCGCCGCTACTAAGGCAATCCCTGTTGGTTTCTTTTCCACCGCTTTTGATATGCAAA |
| *EC102* | F | Aureobasidium_pullulans | Epilobium_canum | CCCGACCTCCAACCCTTTGTTGTTAAAACTACCTTGTTGCTTTGGCGGGACCGCTCGGTCTCGAGCCGCTGGGGATTCGTCCCAGGCGAGCGCCCGCCAGAGTTAAACCAAACTCTTGTTATTTAACCGGTCGTCTGAGTTAAAATTTTGAATAAATCAAAACTTTCAACAACGGATCTCTTGGTTCTCGCATCGATGAAGAACGCAGCGAAATGCGATAAGTAATGTGAATTGCAGAATTCAGTGAATCATCGAATCTTTGAACGCACATTGCGCCCCTTGGTATTCCGAGGGGCATGCCTGTTCGAGCGTCATTACACCACTCAAGCTATGCTTGGTATTGGGCGTCGTCCTTAGTTGGGCGCGCCTTAAAGACCTCGGCGAGGCCACTCCGGCTTTAGGCGTAGTAGAATTTATTCGAACGTCTGTCAAAGGAGAGGAACTCTGCCGACTGAAACCTTTATATTTTTCTAGGTTGACCTCGGATCAGGTAGGGATACCCGCTGAACTT |
| *EC69* | F | Metschnikowia_koreensis | Epilobium_canum | GCTCAAATTTGAAATCCTCCGGGAATTGTAATTTGAAGGTGGGGTTGAATAGGTCTAGATACTTTAAGTCCATTGGAAAATGGCGCCMTGGAGGGTGATAGCCCCGTAAAAGTATTCAAACCTTCTTTTCTTCCCCTCCTAAGARTCGAGTTGTTTGGGAATGCAGCTCTAGTGGGTGGTAAATTCCATCTAAAGCTAAATATTGGCGAGAGACCGATAGCGAACAAGTACAGTGATGGAAAGATGAAAAGCACTTTGAAAAGARAGTGAAAAAGTACGTGAAATTGTTGAAAGGGAAGGGCTTGCAAGCAGACACAACCTCGGTTGGGCCAGCATCGGARTGGGGGGARACAAAAAAGGTTA |
| *SCC187* | B | Pantoea_agglomerans | Calystegia_occidentalis | TCTTTTGCAACCCACTCCCATGGTGTGACGGGCGGTGTGTACAAGGCCCGGGAACGTATTCACCGTGGCATTCTGAGCCAKGATCAMAMTCTSWTCCSACTTCACGGAGTCSAGTTGCAGACTCCGATCCGGACTACGACGCACTTTGTGASGTCCGCTTGCTCTCSCGAGGTCSCTTCTCTTTGTATGCGCCATTGTAACACGTGTGTARCCCTCTCGTAAGGGCCATGATGACTTGACGTCATCCCCACCTTCCTCCGGTTTATCACCGGCAGTCTCCTTTGAGTTCCCGACCGAATCGCTGGCAACAAAGGATAAGGGTTGCGCTCGTTGCGGGACTTAACCCAACATTTCACAACACGAGCTGACGACAG |

| | | | | |
|---|---|---|---|---|
| | | | | CCATGCAGCACCTGTCTCASCGTTCCCGAAGGCA |
| *SCC31* | B | Staphylococcus_warneri | Stachys_ajugloides | CATGCTGATCTACGATTACTAGCGATTCCAGCTTCATGTAGTCGAGTTGCARACTACAATCCRAACTGAGAACAACTTTATGGGATTTGCTTGACCTCGCGGTTTAGCTGCCCTTTGTATTGYCCATTGTAGCACGTGTGTASCCCAAATCATAAGGGGCATGATGATTTGACGTCATCCCCACCTTCCTCCGGTTTGTCACCGGCAGTCAACTTAGAGTGCCCAACTTAATGATGGCAACTAAGCTTAAGGGTTGCGCTCGTTGCGGGACTTAACCCAACATCTCACGACACGAGCTGACGACAACCATGCACCACCTGTCACTTTGTCCCCCGAAGGGGAAGACTCTATCTCTAGAGCGGTCAAAGGATGTCAAGATTTGGTAAGGTTCTTCGCGTTGCTTCRAATTAAACCACATGCTCCACCGCTTGTGCGGGTCCCCGTCAATTCCTTTGAGTTTCAACCTTGCGGTCGTACTCCCCAGGCGGAGTGCTTAATGCGTTAGCTGCAGCACTAAGGGGCGGAAACCCCCTAACACTTAGCACTCATCGTTTACG |
| *Baugh con int N Y1* | F | Cryptococcus_victoriae (Vishniacozyma_victoriae) | Prunus_amygdolus | GAGCTCAAATTTAAAATCTGGCGTCTTTCAGGCGTCCGAGTTGTAATCTATAGAGGCGTTTTCCGCGCCGGACCGCGTCCAAGTCTCTTGGAATAGAGTATCAAAGAGGGTGACAATCCCGTACTTGACGCGACAACCGGTGCTCTGTGATACGTTCTCAATGAGTCGAGTTGTTTGGGAATGCAGCTCTAAATGGGTGGTAAATTCCATCTAAGGCTAAATATTGGCGAGAGACCGATAGCGAACAAGTACCGTGAGGGAAAGATGAAAAGCACTTTGGAAAGAGAGTTAAACAGCACGTGAAATTGTTAAAAGGGAAACGATTGAAGTCAGTCGTGTGAAAGGTATTCAGCCGTCTCTGGCGGTGTATTTGCCTTTCACGGGTCAACATCAGTTTGATCCGGTAGAAAAAGGCAGGAGGAAGGTGGCACCCTCGGGTGTGTTATAGCCTCTTGTCATATGTGCCGGACCAGACTGAGGAACGCAGCTTGCCGCAAGGCCGGGGTTCGCCCACGTACAAGCTTAGGATGTTGACATAATGGCTT |

**Supplementary Table S2** Plant species sampled, their abbreviations, number of individual flowers sampled at each site and nectary location for each plant species.

| **Species abbreviation** | **Plant species** | **Bee Bio** | **Stebbins** | **Nectary location** |
|---|---|---|---|---|
| Aescal | *Aesculus californica* | 0 | 57 | Within_corolla_mid |
| Amsmen | *Amsinckia menziesii var. intermedia* | 8 | 28 | Within_corolla_short |
| Antvex | *Antirrhinum vexillo-calyculatum* | 0 | 27 | Within_corolla_mid |
| Asccor | *Asclepias* | 0 | 15 | Within_corolla_short |
| Asceri | *Asclepias eriocarpa* | 51 | 0 | Within_corolla_short |
| Ascfas | *Asclepias fascicularis* | 14 | 19 | Within_corolla_short |
| Broele | *Brodiaea elegans* | 0 | 6 | Within_corolla_mid |
| Calocc | *Calystegia occidentalis* | 0 | 105 | Within_corolla_short |
| Casocc | *Castilleja spp.* | 0 | 19 | Within_corolla_long |
| Chlpom | *Chlorogalum pomeridianum* | 0 | 32 | Exposed |
| Clacon | *Clarkia concinna* | 0 | 20 | Within_corolla_mid |
| Clapur | *Clarkia purpurea ssp. quadrivulnera* | 2 | 4 | Within_corolla_short |
| Claung | *Clarkia unguiculata* | 36 | 47 | Within_corolla_short |
| Clawil | *Clarkia williamsonii* | 19 | 0 | Within_corolla_short |
| Colhet | *Collinsia heterophylla* | 45 | 71 | Within_corolla_mid |
| Colspa | *Collinsia sparsiflora* | 0 | 6 | Within_corolla_mid |
| Cyngra | *Cynoglossum grande* | 0 | 18 | Within_corolla_mid |
| Delhes | *Delphinium hesperium* | 0 | 70 | Within_corolla_long |
| Delnud | *Delphinium nudicaule* | 0 | 65 | Within_corolla_long |
| Delpat | *Delphinium patens* | 0 | 24 | Within_corolla_long |
| Diccap | *Dichelostemma capitatum* | 0 | 17 | Within_corolla_mid |
| Dicvol | *Dichelostemma volubile* | 0 | 50 | Within_corolla_mid |
| Epican | *Epilobium canum* | 0 | 6 | Within_corolla_long |
| Erical | *Eriodictyon californicum* | 0 | 6 | Within_corolla_short |
| Erysuf | *Erysimum suffrutescens* | 5 | 0 | Within_corolla_short |
| Lamamp | *Lamium amplexicaule* | 0 | 5 | Within_corolla_mid |

| | | | | |
|---|---|---|---|---|
| Lepcal | *Lepechinia calycina* | 0 | 23 | Within_corolla_mid |
| Mimaur | *Mimulus aurantiacus* | 0 | 86 | Within_corolla_long |
| Monvil | *Monardella villosa "coyote mint"* | 48 | 36 | Within_corolla_mid |
| Penhet | *Penstemon heterophyllus* | 63 | 27 | Within_corolla_mid |
| Phacal | *Phacelia californica* | 47 | 43 | Within_corolla_short |
| Phacam | *Phacelia campanularia* | 41 | 0 | Within_corolla_short |
| Phatan | *Phacelia tanacetifolia* | 9 | 0 | Within_corolla_short |
| Salcoa | *Salvia coahuilensis "purple ginny"* | 69 | 0 | Within_corolla_mid |
| Salcol | *Salvia columbariae* | 29 | 0 | Within_corolla_mid |
| Salmic | *Salvia microphylla "hot lips"* | 84 | 0 | Within_corolla_mid |
| Salspa | *Salvia spathacea* | 22 | 0 | Within_corolla_long |
| Scrcal | *Scrophularia californica* | 118 | 93 | Within_corolla_short |
| Staaju | *Stachys ajugoides var. rigida* | 0 | 9 | Within_corolla_mid |
| Toxfre | *Toxicoscordion fremontii* | 0 | 8 | Exposed |
| Trilan | *Trichostema lanatum* | 37 | 0 | Within_corolla_mid |
| Trilax | *Triteleia laxa* | 0 | 42 | Within_corolla_long |

**Supplementary Table S3.** ANOVA tables (type III sums of squares) for best-fit models chosen with AIC from a model containing all 2-way interactions among predictors (Julian date, site, year). Significance of model terms was calculated using LRT and $\chi^2$ statistics for models including microbial presence, and F-tests for models including microbial abundance (log-transformed).

**Table S3A. CFU presence on R2A plates (bacteria)**

| | Df | Deviance | AIC | LRT | Pr(>Chi) | |
|---|---|---|---|---|---|---|
| <none> | | 2403.7 | 2415.7 | | | |
| Julian | 1 | 2458.9 | 2468.9 | 55.188 | 1.10E-13 | *** |
| Site | 1 | 2410.3 | 2420.3 | 6.565 | 0.0104 | * |
| Year | 1 | 2440.6 | 2450.6 | 36.834 | 1.29E-09 | *** |
| Site:Year | 1 | 2410.3 | 2420.3 | 6.562 | 0.01042 | * |
| Julian:Year | 1 | 2458.9 | 2468.9 | 55.198 | 1.09E-13 | *** |

**Table S3B. CFU presence on YMA plates (fungi)**

| | Df | Deviance | AIC | LRT | Pr(>Chi) | |
|---|---|---|---|---|---|---|
| <none> | | 1726.6 | 1738.6 | | | |
| Julian | 1 | 1750.6 | 1760.6 | 23.987 | 9.70E-07 | *** |
| Site | 1 | 1761.6 | 1771.6 | 34.968 | 3.35E-09 | *** |
| Year | 1 | 1742.8 | 1752.8 | 16.166 | 5.80E-05 | *** |
| Julian:Site | 1 | 1752.7 | 1762.7 | 26.054 | 3.32E-07 | *** |
| Julian:Year | 1 | 1750.7 | 1760.7 | 24.013 | 9.57E-07 | *** |

**Table S3C. CFU abundance (bacteria)**

| | Df | SumofSq | RSS | AIC | F | Pr(>F) |
|---|---|---|---|---|---|---|
| <none> | | 4508.5 | 1663.2 | | | |
| Julian | 1 | 238.877 | 4747.4 | 1755.1 | 95.954 | 2.20E-16 |
| Site | 1 | 62.482 | 4571 | 1686.3 | 25.098 | 5.98E-07 |
| Year | 1 | 182.485 | 4691 | 1733.3 | 73.301 | 2.20E-16 |

| | | | | | | |
|---|---|---|---|---|---|---|
| Site:Year | 1 | 62.482 | 4571 | 1686.3 | 25.098 | 5.98E-07 |
| Julian:Year | 1 | 238.937 | 4747.4 | 1755.1 | 95.978 | 2.20E-16 |

**Table S3D. CFU abundance (fungi)**

| | Df | SumofSq | RSS | AIC | F | Pr(>F) |
|---|---|---|---|---|---|---|
| <none> | | 1073.8 | -939.16 | | | |
| Julian | 1 | 17.417 | 1091.2 | -911.97 | 29.326 | 6.94E-08 |
| Site | 1 | 21.346 | 1095.1 | -905.45 | 35.941 | 2.45E-09 |
| Year | 1 | 10.92 | 1084.7 | -922.8 | 18.387 | 1.90E-05 |
| Julian:Site | 1 | 15.053 | 1088.8 | -915.9 | 25.347 | 5.27E-07 |
| Julian:Year | 1 | 17.43 | 1091.2 | -911.95 | 29.348 | 6.86E-08 |

**Supplementary Table S4**.

Microbial genera identified from floral nectar samples from a total of 309 colonies analyzed using MALDI and Biotyper (see supplementary methods S1). Colonies were sampled haphazardly from the collection. Cells list occurrences by flower morphology, along with the total number. Filamentous fungi could not be identified using this method and the bacterial genus *Gluconobacter* (*Neokomagataea*) is only inconsistently identified using this method. As a result, *Neokomagataea* colonies are likely to be included in the 'no spectra or no match' category, likely along with other taxa not in our database.

| Genus | Kingdom | Exposed | Short corolla | Mid corolla | Long corolla | Total |
|---|---|---|---|---|---|---|
| *Metschnikowia* | Fungi | 0 | 7 | 24 | 35 | 66 |
| *Acinetobacter* | Bacteria | 0 | 23 | 12 | 2 | 37 |
| *Bacillus* | Bacteria | 0 | 6 | 19 | 5 | 30 |
| *Pseudomonas* | Bacteria | 0 | 3 | 3 | 11 | 17 |
| *Erwinia* | Bacteria | 0 | 5 | 7 | 1 | 13 |
| *Candida* | Fungi | 1 | 6 | 4 | 0 | 11 |
| *Lactobacillus* | Bacteria | 1 | 1 | 6 | 0 | 8 |
| *Micrococcus* | Bacteria | 1 | 4 | 1 | 2 | 8 |
| *Staphylococcus* | Bacteria | 0 | 5 | 3 | 0 | 8 |
| *Pantoea* | Bacteria | 0 | 4 | 0 | 2 | 6 |
| *Rosenbergiella* | Bacteria | 0 | 0 | 0 | 5 | 5 |
| *Clostridium* | Bacteria | 0 | 3 | 1 | 0 | 4 |
| *Salmonella* | Bacteria | 0 | 1 | 2 | 1 | 4 |
| *Arthrobacter* | Bacteria | 0 | 2 | 1 | 0 | 3 |
| *Brenneria* | Bacteria | 0 | 0 | 0 | 3 | 3 |
| *Citrobacter* | Bacteria | 0 | 1 | 0 | 2 | 3 |
| *Filifactor* | Bacteria | 0 | 0 | 1 | 2 | 3 |
| *Aureobasidium* | Fungi | 0 | 1 | 0 | 1 | 2 |
| *Blautia* | Bacteria | 0 | 0 | 1 | 1 | 2 |
| *Escherichia* | Bacteria | 0 | 0 | 0 | 2 | 2 |
| *Listeria* | Bacteria | 0 | 1 | 1 | 0 | 2 |

| | | | | | | |
|---|---|---|---|---|---|---|
| *Moraxella* | Bacteria | 0 | 1 | 1 | 0 | 2 |
| *Streptomyces* | Bacteria | 0 | 1 | 1 | 0 | 2 |
| *Terrimonas* | Bacteria | 0 | 1 | 0 | 1 | 2 |
| *Agromyces* | Bacteria | 0 | 1 | 0 | 0 | 1 |
| *Arsenophonus* | Bacteria | 0 | 0 | 1 | 0 | 1 |
| *Austwickia* | Bacteria | 0 | 0 | 0 | 1 | 1 |
| *Brevibacillus* | Bacteria | 0 | 1 | 0 | 0 | 1 |
| *Brevibacterium* | Bacteria | 0 | 0 | 1 | 0 | 1 |
| *Capnocytophaga* | Bacteria | 0 | 0 | 1 | 0 | 1 |
| *Corynebacterium* | Bacteria | 0 | 1 | 0 | 0 | 1 |
| *Cryptococcus* | Fungi | 0 | 0 | 0 | 1 | 1 |
| *Curtobacterium* | Bacteria | 0 | 0 | 1 | 0 | 1 |
| *Enterobacter* | Bacteria | 0 | 0 | 0 | 1 | 1 |
| *Marinibacillus* | Bacteria | 0 | 0 | 0 | 1 | 1 |
| *Mycoplasma* | Bacteria | 0 | 0 | 1 | 0 | 1 |
| *Nocardia* | Bacteria | 0 | 1 | 0 | 0 | 1 |
| *Paenibacillus* | Bacteria | 0 | 1 | 0 | 0 | 1 |
| *Pichia* | Fungi | 0 | 1 | 0 | 0 | 1 |
| *Rahnella* | Bacteria | 0 | 0 | 0 | 1 | 1 |
| *Rhizobium* | Bacteria | 0 | 1 | 0 | 0 | 1 |
| *Solibacillus* | Bacteria | 0 | 0 | 1 | 0 | 1 |
| *Sphingobacterium* | Bacteria | 0 | 0 | 0 | 1 | 1 |
| *Sphingobium* | Bacteria | 0 | 0 | 1 | 0 | 1 |
| *Sphingomonas* | Bacteria | 0 | 1 | 0 | 0 | 1 |
| *Streptococcus* | Bacteria | 0 | 1 | 0 | 0 | 1 |
| No match | | 1 | 18 | 17 | 8 | 44 |

**Supplementary Methods S1**

*Lab dispersal assay: colony maintenance and assay setup*

We started a colony of western flower thrips, *Frankliniella occidentalis*, with thrips obtained from a lab colony held by Diane Ullman at UC Davis since 1995, and originating from the Kamilo Iki valley on the island of O‘ahu, Hawaii (Bautista et al. 1995). We reared thrips on green bean pods (*Phaseolus vulgarisas*) as previously described (Ullman et al. 1992). Dispersal ability via thrips activity was estimated by counting the number of previously unoccupied wells that became occupied by microbes after 24 hours of thrips activity by 5 newly eclosed adult female thrips. The landscape layout consisted of 64 unoccupied wells, each with 200uL of sterile TSB (tryptic soy broth, with added sucrose and glucose 75 g/L) and 32 occupied wells of a single strain of microbe beings assayed (200uL at 10,000 cells/uL in TSB). Columns 2, 5, 8, and 11 were occupied by microbes, resulting in every unoccupied well on the plate having one direct neighbor containing the microbe of interest. Control plates contained 5 thrips, with all 96 wells containing 200uL of sterile TSB.

After 24 hours for foraging at 30 °C, thrips were killed by pipetting 16 separate 50uL drops of ethyl acetate into the well plate. Microbes were cultured for an additional 5 days, and each well was measured for optical density (OD) at 600nm. Wells with OD substantially higher (mean plus 6 standard deviations) than a control plate with no thrips or microbes were determined to be occupied. Calibration plates with serial dilutions confirmed positive growth rates during all trials.

Sterilization of thrips was attempted by feeding the thrips sterile TSB with Rifampicin (0.2mg/mL) and cycloheximide (0.4 mg/mL) for 48 hours and then vortexing thrips in 30% ethanol (5s), sterile water (15s), 5% bleach (90s), sterile water (15s), sterile water (15s), and sterile water (15s). Thrips were vortexed in wash solutions in cohorts of approximately 50 using “wash chambers”, consisting of 1000uL barrier pipette tips with added no-thrips mesh to contain multiple thrips at once and allow plunging of wash solutions. After the final wash, thrips were centrifuged at 1000 rpm for 10s to absorb remaining moisture. Contaminants persisted on thrips at low frequencies following this sterilization procedure but were low enough in abundances such that control plates had consistently low numbers of occupied wells.

MALDI-TOF confirmation of microbe identity was performed during a subset of trials to confirm microbial identity of dispersed microbes, with contaminants appearing on treatment plates with a frequency that matched contamination levels seen in controls.

We isolated floral microbes from flower nectar from naturally occurring flowers and planted cultivars, bee brood cells, as well as thrips mouthparts to capture taxonomic diversity of common nectar microbes.

Microbes included in the assay were isolated from floral nectar, bee digestive tracts, and from thrips mouthparts. Microbes were chosen to represent several of the most common strains from collections in this study and are summarized in Supplemental Table 4 along with Sanger sequencing results for 16S and NL1 regions of the mitochondria, host species information, and isolate specific strain names.

Ullman, D. E. et al. 1992. A midgut barrier to Tomato spotted wilt virus acquisition by adult western flower thrips. – Phytopathology 82: 1333 – 1342.

**Supplementary Methods S2**

*Correlation between cell and CFU counts*

We grew out *Metschnikowia reukaufii* and *Acinetobacter nectaris* from freezer stock onto yeast media agar (YMA) containing chloramphenicol and Reasoner's agar containing cycloheximide (with 16% sucrose). After 3 days of growth at 25°C, colonies of each microbe were suspended in phosphate buffered saline. Each microbial suspension was vortexed for 90 seconds to break up cell aggregates and pipetted onto a hemocytometer where the cell density was counted under a microscope. Each microbial suspension was then serially diluted (dilution amounts listed below) and 100ul of each dilution was plated onto YMA and R2A antimicrobial supplemental plates (media recipes as above), and incubated for 3 days after which the colony forming units on each plate were counted.

The assay was performed over 3 different trials with YM and R2A each tested twice using new growth from freezer stock and new cell counts and dilutions to ensure consistency in the results. On trial 1, yeast counts only were performed at 50 cells/ul, 10 cells/ul, 1 cell/ul, 0.1 cells/ul and 0.01 cells/ul (n=3). On trial 2, bacteria counts only were performed at 100 cells/ul, 10 cells/ul, 1 cell/ul (n=4). On trial 3 both yeast and bacteria counts were performed at 50 cells/ul, 10 cells/ul, 1 cell/ul, 0.1 cells/ul and 0.01 cells/ul (n=4).

A linear model was constructed with a log base 10 transformation of the number of cells plated as the predictor variable and log base 10 transformation of CFUs counted plus one as the response.

**Supplementary Methods S3**

*Microbial culture identification*

We identified a subset of microbial colonies from nectar samples. Bacterial and yeast isolates from glycerol stocks were plated on isolation media, with isolates chosen haphazardly from early, mid and late in the flowering season. Material from a single representative colony for each isolate was spotted in duplicate onto a MTP 384 Ground Steel Target Plate (Bruker Daltonics), overlaid with 1uL 70% formic acid, and allowed to air dry. 1uL of alpha-cyano-4-hydroxycinnamic acid (HCCA) matrix dissolved in a 50:45:5 solution of acetonitrile, water, and trifluoroacetic acid was added and let dry [same prep as in Seuylemezian 2018, JPL microbes paper. doi: 10.3389/fmicb.2018.00780]. Spectra were obtained using an ultrafleXtreme MALDI-TOF instrument (Bruker Daltonics, Billerica, MA, United States). Spectra for each isolate were then compared with a custom in-house library of main spectral profiles (MSPs) and the Bruker libraries (Bacteria, Eukaryotes) using MBT Compass Explorer software (Bruker Daltonics, Billerica, MA, United States). Log scores between 1 and 3 were generated for each isolate’s top 10 best matches. Scores ≥2.0 are considered close matches; lower scores are likely species not present in the library but genus identification for those with matches above 1.6 were used. Matches for each isolate were compiled and compared across sampling timepoints and between sites.
